# Investigator-blind discovery of structural elements controlling GPCR function

**DOI:** 10.64898/2026.03.22.713462

**Authors:** Jingjing Ji, Edward Lyman

## Abstract

With the advance of hardware and software for molecular dynamics simulation it has become routine to obtain trajectories that are tens of microseconds in duration for all kinds of protein machinery. This shifts the burden of work onto analysis of the simulation data and opens opportunities for more rigorous and reproducible observations on mechanism. Toward this end we developed an investigator-blind analysis pipeline which operates on featurized simulation data, performs unsupervised clustering, and then identifies which input features are most discriminatory of cluster identity. Application of this pipeline to a large set of G-protein coupled receptor simulation data shows that it identifies several well-known microswitches. Inspection of these structural elements reveals changes in conformation that are known to accompany functional transitions of the receptor. In addition to these known structural elements the analysis also identifies two possibly new structural motifs: the kink in transmembrane helix 2, and a coupled “piston-like” motion of TM2 and TM3.

## Introduction

The G-protein coupled receptors transduce an incredible diversity of chemical signals across the cell membranes of metazoans. At the time of writing more than 200 individual GPCR structures have been determined (*1*), including members from all classes (A through F), bound to ligands of all different kinds of activity, and including examples of GPCRs coupled to heterotrimeric G-proteins or to arrestins. These structural data have provided invaluable insight into the mechanisms by which ligands influence signaling, and how signaling partners couple to the intracellular face of the receptor (*2–5*).

These structural data have also enabled a massive amount of simulation data to be collected by groups from around the world. At the time of writing the GPCRmd repository (*6*) holds more than 600 simulations totaling over 2 msec of simulation time. In analyzing simulation data, one often wishes to identify different conformational states that are sampled during the simulation, in order to assess structure-function relationships. This requires first projecting the 3N Cartesian coordinates of the N atoms of the protein into a reduced dimensionality space, then identifying sets of similar conformations by a (typically unsupervised) clustering method.

The biomolecular simulation field has been grappling with these problems for decades. Principle component analysis (PCA) was first applied to protein simulation data in 1992 by Garcia (*7*), and shortly thereafter by Berendsen (*8*). In these early applications, the “features” of the protein were simply taken to be the x,y,z coordinates of the atoms of the protein. In this representation, the eigenvectors are linear combinations of atom displacements, and naturally describe (sometimes functional) collective motions of the protein. For this reason, Berendsen and coworkers called this approach “essential dynamics.” A related approach called time-lagged independent component analysis (tICA) (*9*, *10*) incorporates a time lag in order to identify the slowest PCA dimensions, and is widely used in the construction of Markov models.

Many groups have exploited recent developments in machine learning to make new progress on this old problem (*11–15*). There are now several algorithms for dimensionality reduction, such as t-distributed stochastic neighbor embedding (t-SNE) (*16*), multi-dimensional scaling (MDS) (*17*), autoencoders (*18*, *19*), and uniform manifold approximation and projection (UMAP) (*20*, *21*). For the unsupervised learning/clustering step there are also many options to choose from, including k-means clustering, spectral clustering, Gaussian Mixture Models, Density-Based Spatial Clustering of Applications with Noise (DBSCAN) (*22*), and hierarchical DBSCAN (HDBSCAN) (*23*). Indeed, upon entering this arena one may feel paralyzed by the algorithmic choices (and their combinations).

Implementations for all are available either in Scikit-learn https://scikit-learn.org/stable/ or from GitHub repositories, lowering the barrier to trying different approaches on one’s problem of the moment. We therefore decided to try several different commonly used approaches in order to assess their strengths and weaknesses for mapping the conformational landscape of a GPCR. We assembled a simulation dataset for the A_2A_ adenosine receptor totalling nine different trajectories, ranging in duration from 1 to 10 *μ*sec. The simulations were initiated from different states — apo, or bound to ligands of differing efficacy, with and without a bound G-protein, and in a variety of different membrane environments. By assembling such a diverse set of trajectories we hoped to sample as broadly as possible the conformational landscape of the protein, and perhaps observe transitions between different substates relevant for function.

We find that the combination of UMAP for dimensionality reduction and HDBSCAN for clustering is well-suited to the analysis of GPCR simulation data, as it produces clusters which correspond to physically meaningful receptor states. This is shown by projecting the trajectory data (featurized by inverse alpha carbon distances) onto selected UMAP dimensions, and analyzing the clusters identified by HDBSCAN. A significant advantage of UMAP (unlike t-SNE) is that it preserves distances in the latent space, so that clusters that are near to each other in the latent space represent physically similar conformations. Thus, simulations which are initiated from similar conformations but under different conditions (lipid environment, ligation state) initially remain close together in the projected space, before exploring new states. This is illustrated below by considering several simulation trajectories which all begin from the structure of the fully active receptor coupled to an engineered G-protein. When simulated with both G-protein and agonist bound the simulation remains within a single cluster. But when the initial state is perturbed (by deleting the G-protein, for example) the receptor relaxes into new region of conformation space, identified by HDBSCAN as a new cluster. Features that discriminate the clusters were then found by training a classifier (XGBOOST) to predict cluster identity from the features and then interpreting the classifier with SHAP. This analysis workflow frequently identifies already described activation motifs or “microswitches” — NPxxY, D/ERY, PIF motifs (*2*, *24*, *25*). For example, after deletion of the G-protein, the classifier identifies the NPxxY motif as a key feature discriminating the relaxed from the fully active cluster. Upon inspection it is observed to “untwist,” shifting the receptor toward an inactive state.

Many other examples of this type are described in the following and discussed in the context of published models for receptor activation. The workflow also identifies at least two (to our knowledge) new structural features or microswitches: a straightening of the kink in TM2 (near the conserved sodium ion binding site), and a coupled motion of TM2 and TM3. Taken together, these results support analysis workflows for MD simulation in which investigator expectations take a back seat to features discovered by automated analysis, followed by human analysis and validation.

## Results

### Data-driven analysis reveals conformational ensembles

We collected a dataset of A_2A_R conformations sampled across nine simulation trajectories initiated from different states, and ranging in duration from 1 to 10 usec (**Table 1**), totalling 28,400 configurations. The initial structures were chosen to represent a range of functional states: bound to antagonist, bound to agonist, or bound to both agonist and an engineered mini G-protein. We also generated from these several other initial structures in which ligands and/or G-protein were deleted, in order to observe how the receptor relaxes following unbinding of ligands or G-protein.

**Table 1.** Simulated systems and conditions.

| System index | Initial state | PDB ID | Ligation state | G protein bound | Bilayer | Duration [μs] | Cluster label |
| --- | --- | --- | --- | --- | --- | --- | --- |
| 1 | active | 5G53 | NECA | Yes | asymmetric MM | 2 | a |
| 2 | active | 5G53 | NECA | No | asymmetric MM | 10 | a, i |
| 3 | active | 5G53 | apo | No | asymmetric MM | 2 | a, f |
| 4 | active | 5G53 | NECA | No | POPC | 4.4 | a, g, h |
| 5 | intermediate | 4UHR | CGS21680 | No | asymmetric MM | 1 | b |
| 6 | intermediate | 4UHR | apo | No | asymmetric MM | 5 | c |
| 7 | inactive | 5UIG | TC | No | asymmetric MM | 1 | j |
| 8 | inactive | 5UIG | apo | No | asymmetric MM | 2 | d |
| 9 | inactive | 3PWH | ZM241385 | No | asymmetric MM | 1 | e |

Each configuration was featurized as a matrix of inverse alpha carbon distances (see Methods) to more heavily weight short range contacts (*12*, *26*), with 15,916 features per configuration. (Only alpha carbons within the transmembrane bundle were included.) We then looked for a low dimensional representation of the dataset, reasoning that subsequent clustering to identify conformational substates would be both more interpretable and more meaningful in a drastically reduced dimensionality space. We tested principal component analysis (PCA, **fig. S1**), t-distributed stochastic neighbor embedding (t-SNE, **fig. S2**), and uniform manifold approximation and projection (UMAP). We found that UMAP offered a compact (low dimension) representation of the space compared to the other methods (see Supplemental Results) and so continued our analysis based on this representation. The conformations projected into the UMAP space were then clustered with “hierarchical density-based spatial clustering of applications with noise” (HDBSCAN). (See Supplemental Methods for details on the hyperparameter sweep for UMAP and HDBSCAN, **Table S1**)

**Fig. 1** shows the ten clusters identified by HDBSCAN (labeled a through j) across nine system trajectories (numbered 1 through 9) based on the UMAP embedding, projected onto two UMAP dimensions (out of nine total dimensions). Although some of the clusters appear to overlap in this projection, when viewed in three UMAP dimensions they are clearly distinct (see **Supplemental Movie S1**). Each cluster is an ensemble of configurations that are close to one another in the space of alpha-carbon pairwise distances.

**Fig. 1.**
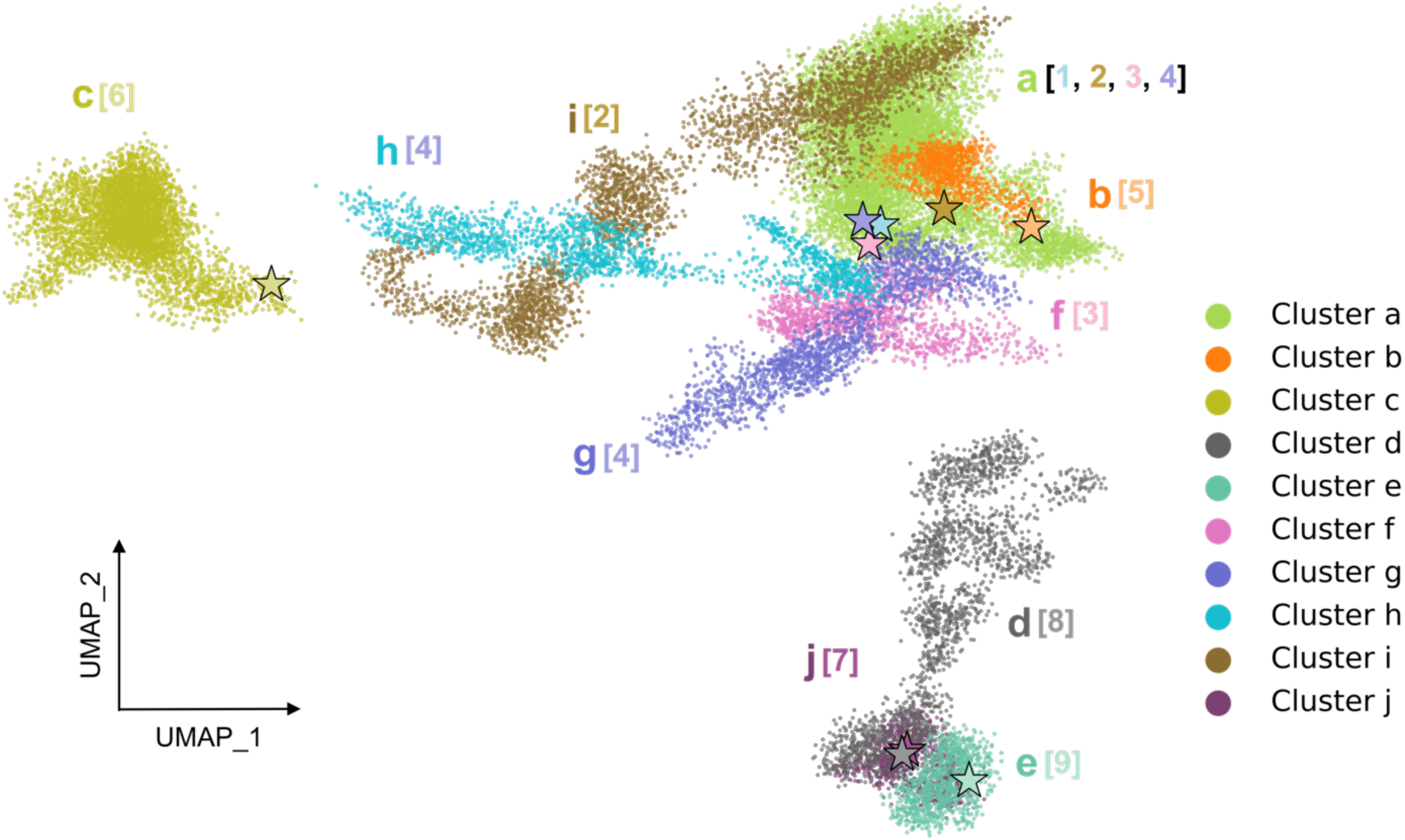
HDBSCAN clustering on the UMAP projection uncovers distinct conformational clusters (labeled as a-j) across nine simulated systems. System indices (in brackets) correspond to Table 1. The stars denote the initial conformations of the nine systems used in the production runs.

The initial structures for systems 1 through 4 were all based on the fully active, G protein-coupled structure (**Table 1**). System 1 was simulated with both the agonist NECA and the mini G-protein bound. The initial structure for system 2 was the same as system 1, except that the G-protein was deleted; for system 3 both the G-protein and NECA were deleted. System 4 was initiated from the same structure as system 2 (with NECA but no G-protein), but was embedded in a different membrane environment. We anticipated that by comparing the conformational space sampled by these systems we might learn how the fully active receptor relaxes after unbinding of ligand and/or G-protein.

The receptor with both G-protein and NECA bound (system 1) remains locked in a fully active state and samples a single cluster, labeled “a”. The other three systems initiated from the fully active state all begin in the same cluster, but then relax into new and distinct regions of conformation space (**Fig. 1**): Without the G protein, the simulation samples configurations that populate cluster i, with neither the G protein nor the ligand the simulation populates cluster f, and when solvated by a simple POPC bilayer instead of an asymmetric membrane it sequentially populates clusters g and then h. In the following sections we present the structural features that distinguish these clusters, but first describe the remaining simulations.

The remaining five trajectories are each classified into their own cluster (**Fig. 1** and **fig. S3**): These are systems 5, 6, 7, 8, and 9 (**Table 1**), which are initiated from three different structures of the protein. System 5 is initiated from a structure that is bound to an agonist (CGS21680), and has several mutations designed to stabilize the active state, but was crystallized without a G-protein mimic. The structure of the intracellular face is described in the literature as intermediate-active or partially active (*27*). This system samples a region of conformation space labeled as cluster b. In system 6, the ligand CGS21680 was deleted to create an apo structure, which samples conformations corresponding to cluster c. Systems 7 and 9 are both initiated from inactive states, bound to the antagonists triazole-carboximidamide (TC) and ZM241385, respectively. Both systems sample a very localized neighborhood of conformations corresponding to clusters j and e, respectively, consistent with the inactive receptor having limited conformational flexibility (*3*, *28*). Although HDBSCAN identified these simulations as sampling distinct clusters, these are close to one another in the UMAP projection, indicating structural similarity. (UMAP preserves distances in the low dimensional projection, which is a major strength for this application.) System 8 was simulated in the apo form of system 7 by removing the antagonist TC. Although its initial structure is close to system 7, system 8 diffuses away from system 7 as the simulation progresses, sampling a conformational ensemble identified as cluster d.

### Identifying structural features that discriminate conformational ensembles

We next sought to identify structural features that discriminate the clusters identified in **Fig. 1**, so that they might be understood in the context of established mechanisms of GPCR structure-function. To identify these features we trained a classifier (XGBoost (*29*)) to predict the cluster label for each simulation snapshot from the inverse alpha carbon distances, then used an interpretability method (SHAP analysis (*30*)) to determine which inverse pair distances were most informative of cluster identity. Up to this point (UMAP → HDBSCAN → XGBoost → SHAP analysis) our workflow is entirely blind to investigator bias. Inspection of these features revealed that they report well known structural changes (e.g., outward motion of TM6 upon activation) and localize to previously reported microswitches (D/ERY ionic lock, NPxxY, PIF, CWxP motifs). In two other cases the analysis indicates that prolines in TM2 and TM5 are involved in transitions between conformational substates.

### Structural differences between inactive, intermediate, and fully active states

We used our investigator-blind and automated pipeline to assess structural differences that discriminate between the inactive, intermediate, and fully active conformational ensembles. We reasoned that since much is already known about these structural differences that this comparison would provide a good test of our approach. To this end we used the SHAP/XGBoost analysis to identify features that discriminate cluster j (inactive state bound to ZM241385) from cluster b (intermediate state bound to CGS21680 but no G-protein), and also cluster b from cluster a (fully active state bound to both NECA and G-protein). Classification performance was nearly perfect (**Tables S2-S3** and **fig. S4**).

The intermediate (cluster b) and inactive (cluster j) conformations are discriminated primarily by the displacement of TM6, consistent with prior observations based on structures of active and inactive class A GPCRs (*28*, *31*, *32*). The SHAP analysis identifies the outward displacement of TM6 upon activation by distances between the residues on the intracellular end of TM6 and TM2 (**Fig. 2B panel I** and **fig. S5A panel I**), highlighted particularly by the S47^2.45^–V239^6.41^ distance (**Fig. 2C panel I** and **fig. S6**) (Residues are labeled by their residue number and Ballesteros–Weinstein numbering (*33*) indicated in superscript). Although less significant than the TM2-TM6 distances, the SHAP analysis also detects changes associated with the NPxxY motif on TM7 (**Fig. 2C panel II**). Changes to the NPxxY motif are more significant when comparing the intermediate and fully active states, described next.

**Fig. 2.**
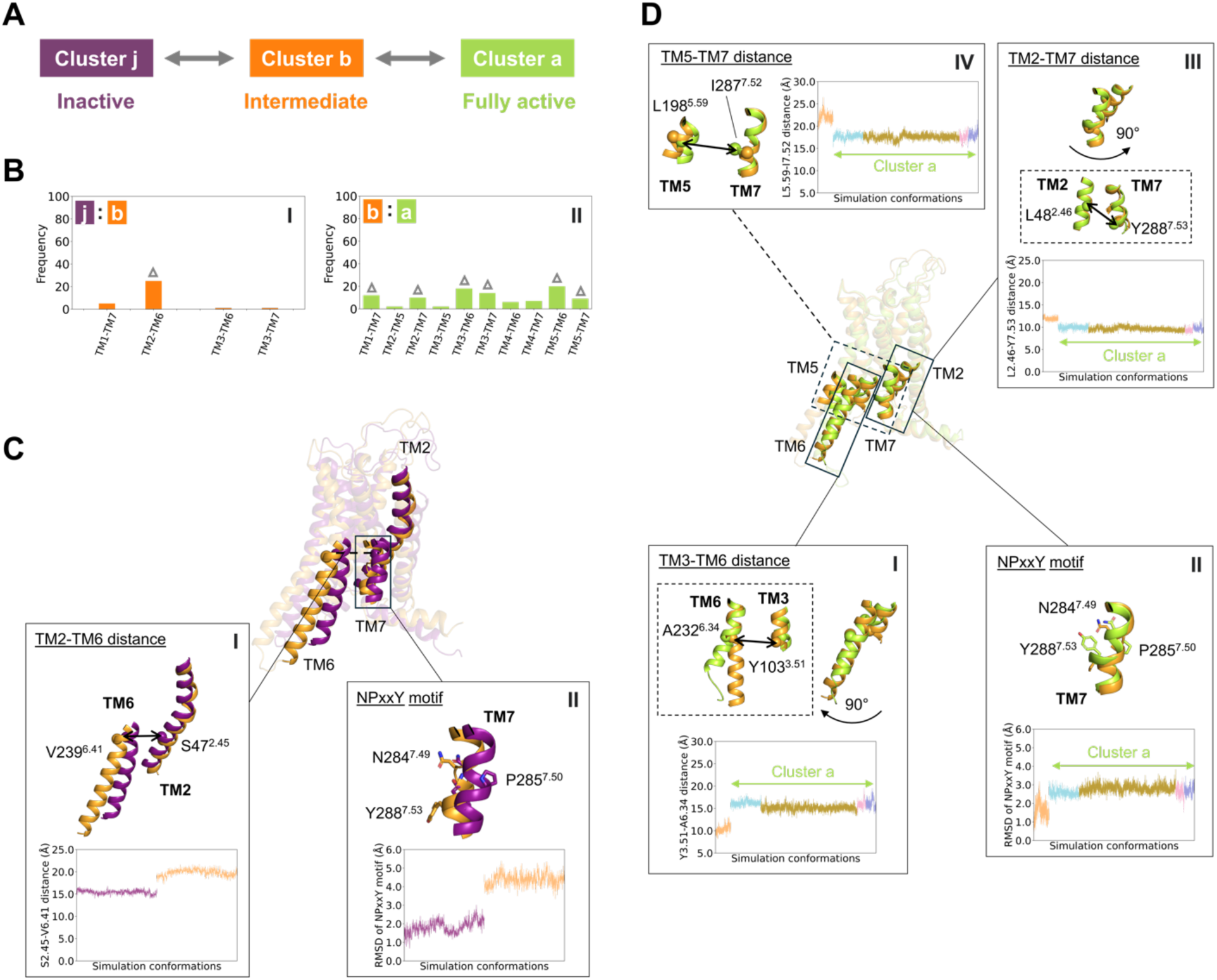
Key conformational differences between clusters. (**A**) Schematic representation of the cluster pairs analyzed. (**B**) Frequencies of transmembrane (TM) pairs in distinguishing between the two clusters. (**C-D**) Key features in the insets distinguishing between the two clusters: (**C**) clusters j and b, (**D**) clusters b and a. Overlaid receptor conformations in (**C**) correspond to the final frames of systems 5 and 7; in (**D**) the final frames of systems 1 and 5. The receptor cartoon conformations are colored by cluster assignments, and the RMSD and residue pair Cα–Cα distance profiles are colored according to their corresponding systems, as shown in panel (A). Cα atoms are shown as spheres, and side chains are shown as sticks.

The residue pairs identified by XGBoost/SHAP identify two well-known structural features that distinguish the fully active conformations (cluster a) from the intermediate conformations (cluster b): the inward movement of the TM6 intracellular end and the untwisting of the NPxxY motif. Although significant features are identified between many pairs of helices (**Fig. 2B panel II**), the largest number of these are localized to TM3–TM6 and TM5–TM6, caused by the inward motion of the intracellular end of TM6. Some of the very highest SHAP values involve residues of the D/ERY motif, particularly the Y103^3.51^–A232^6.34^ distance (**Fig. 2D panel I**). In parallel, the intracellular end of TM5 shifts outward from the helical bundle, leading to variable TM5–TM6 residue pair distances (**fig. S5B**). The contributions of TM3–TM6 and TM5–TM6 residue pairs in distinguishing clusters a and b are shown in **fig. S7**. Furthermore, the conserved NPxxY motif in TM7 “untwists” and shifts outward in cluster b when compared to cluster a, as reflected by its RMSD (**Fig. 2D panel II**). Untwisting of this motif induces changes in alpha carbon distances of L48^2.46^–Y288^7.53^ (TM2–TM7) and L198^5.59^–I287^7.52^ (TM5–TM7) (**Fig. 2D panels III-IV**). Comparable A^2.47^–C^7.54^ and Y^5.58^–Y^7.53^ (Y–Y motif) distances were used as key features in β₂AR deactivation studies (*34*), involving nearby residues to those observed in the present work.

### Structural changes upon ligand unbinding

Systems 5 and 6 are both initiated from an intermediate conformation complexed with an agonist CGS21680; however, in system 6, the ligand was removed to generate the apo form. As the two simulations progress, the two systems sample distinct conformational ensembles labeled b and c respectively in **Fig. 1**. The XGBoost classifier was trained to distinguish these two ensembles. Classification performance was nearly perfect, with results from each of five cross-validation folds presented in **Table S4** and **fig. S8**.

Without the ligand CGS21680 bound, the extracellular portion of the helical bundle exhibits an outward tilt of the extracellular ends of TM2, TM6, and TM7 in cluster c as shown in **Fig. 3C panel I**. This motion is identified by the SHAP analysis through features involving TM2, particularly TM2–TM3, TM2-TM6, and TM2-TM7 residue pair distances (**Fig. 3B** and **fig. S9A**). In recognizing the displacement of TM2 the XGBoost/SHAP analysis zeroes in on several residues that are directly involved with triazole agonist binding (**fig. S10**). On TM2-TM3, distances involving V84^3.32^ and T88^3.36^ increase by about 50% (**Fig. 3C panel II** and **fig. S9B panels I-II**). The “Trp toggle switch” (W246^6.48^) on TM6 moves away from TM2 by a similar amount (**Fig. 3C panel III**), as does S277^7.42^ on TM7 (**Fig. 3C panel IV**). The contributions of additional TM2–TM3, TM2–TM6, and TM2–TM7 residue pairs distinguishing clusters b and c are presented in **fig. S10**. The analysis also identifies an inward displacement of the intracellular end of TM6, toward a more inactive-like conformation, as identified by residue pairs on TM3 and TM6 (**Fig. 3C panel V**).

**Fig. 3.**
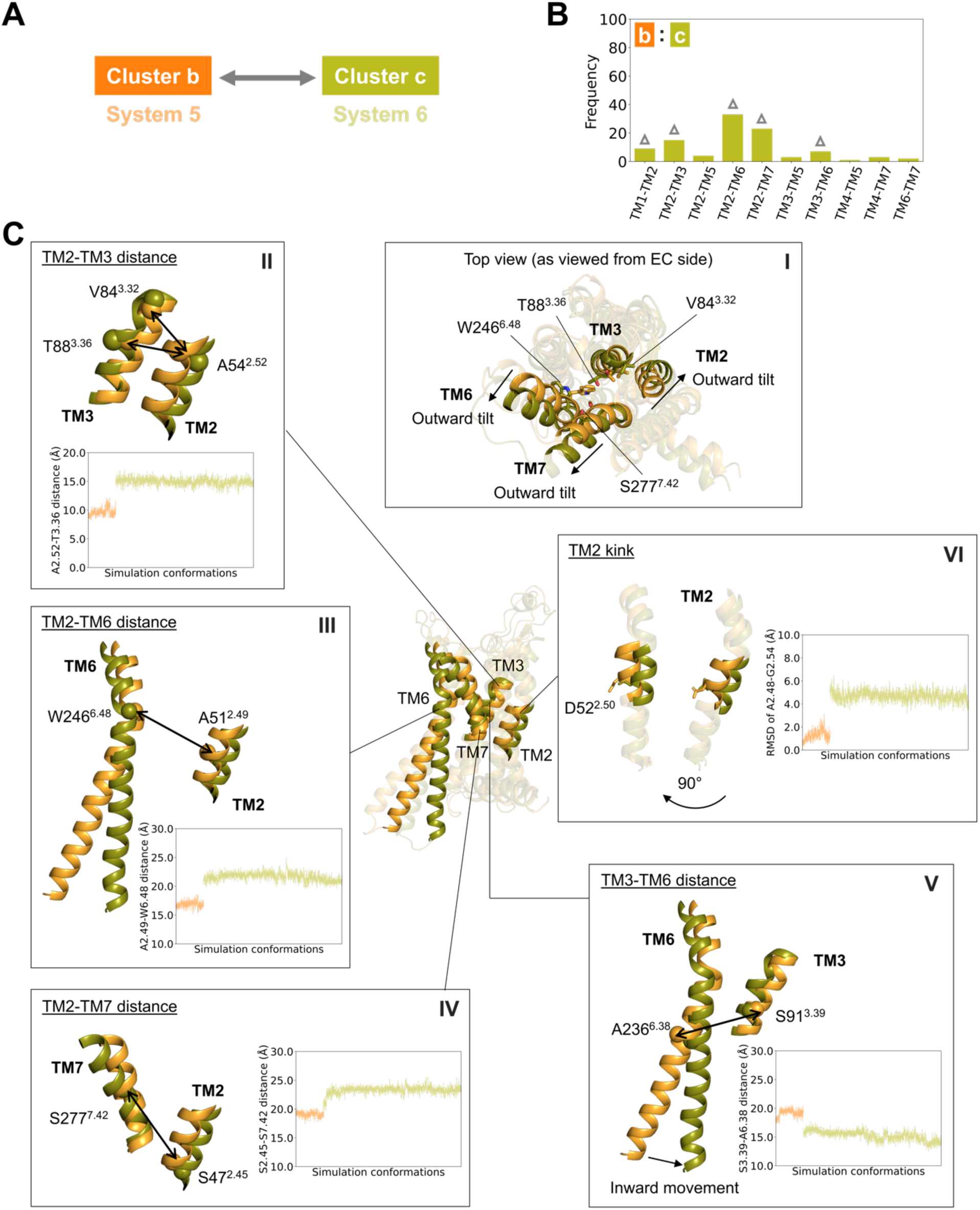
Key conformational differences between the two clusters. (**A**) Schematic representation of the cluster pair analyzed. (**B**) Frequencies of transmembrane (TM) pairs in distinguishing between the two clusters. (**C**) Key features in the insets distinguishing between clusters b and c. Overlaid receptor conformations correspond to the final frames of systems 5 and 6. The receptor cartoon conformations are colored by cluster assignments, and the RMSD and residue pair Cα–Cα distance profiles are colored according to their corresponding systems, as shown in panel (A). Cα atoms are shown as spheres, and side chains are shown as sticks.

The XGBoost/SHAP analysis also identifies an unexpected feature involving the TM2 kink at P61^2.59^. Many of the most highly ranked features involve residues just to the extracellular side of P61^2.59^ (**fig. S10**); inspection of the trajectory and calculation of the RMSD of the TM2 kink shows that it straightens in the apo form compared to the agonist bound intermediate, accompanied by a deviation of the TM2 helical axis (**Fig. 3C panel VI**). As discussed in the next section, a similar transition is observed during relaxation of the fully active (G-protein coupled) receptor state.

### Structural transitions during relaxation of the fully active receptor

Systems 1, 2, 3, and 4 were all initiated from the fully active, G-protein coupled receptor state, but then simulated under different conditions, whereupon they sampled distinct clusters (**Fig. 4A**). We used the XGBoost and SHAP analysis to identify the structural elements that discriminate among these clusters. Classification performance was again nearly perfect, with results from each of five cross-validation folds presented in **Table S5-S8** and **fig. S11**.

**Fig. 4.**
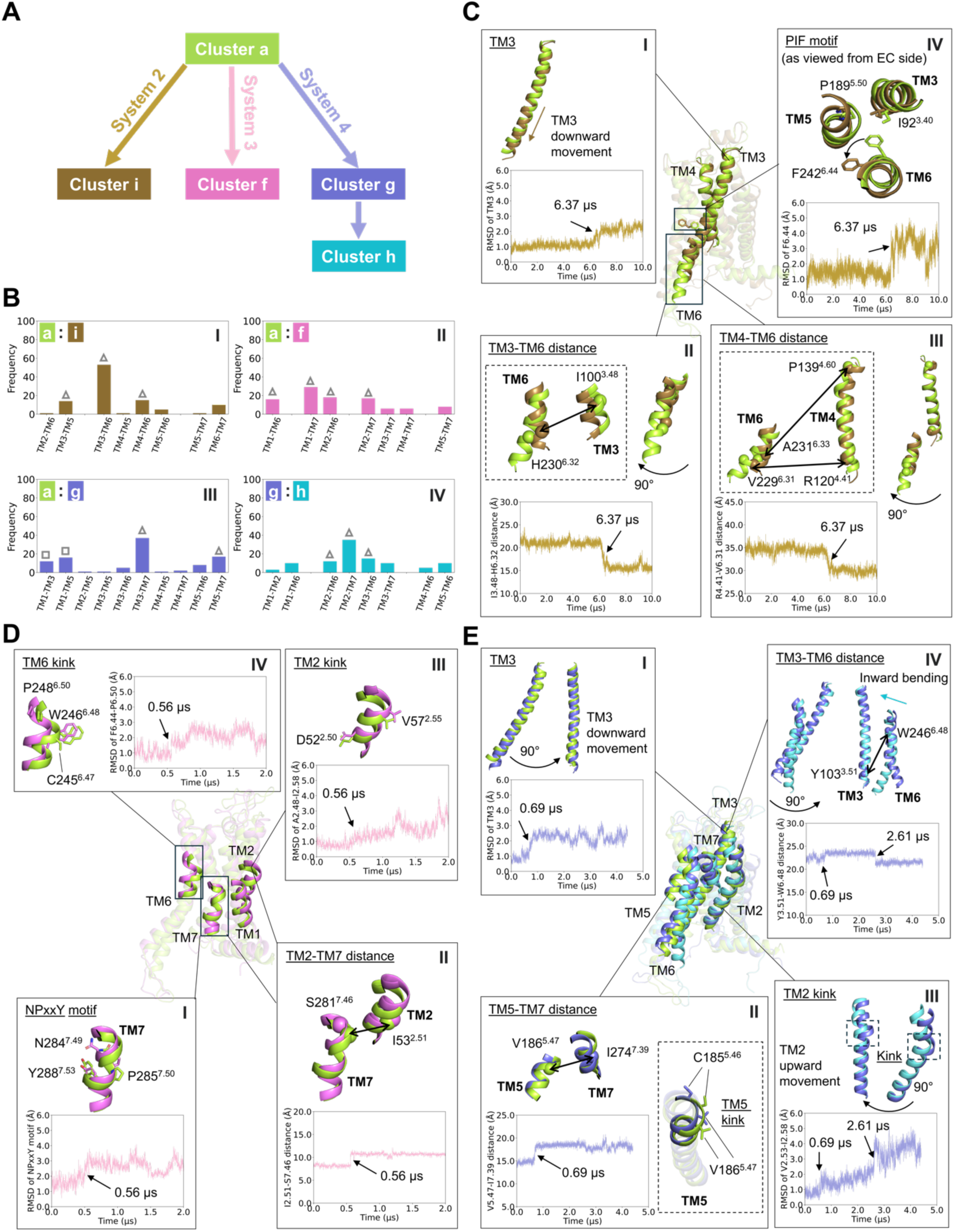
Key conformational changes during simulated state transitions. (**A**) Schematic representation of the state transition observed in simulations. (**B**) Frequencies of transmembrane (TM) pairs in distinguishing between the two clusters. (**C-E**) Key feature changes in the insets associated with state transitions in simulations: (**C**) transition from cluster a to i in system 2, (**D**) transition from cluster a to f in system 3, and (**E**) sequential transitions from cluster a to g and g to h in system 4. Overlaid receptor conformations in (**C**) correspond to the initial and final frames of system 2; in (**D**), the initial and final frames of system 3; and in (**E**), the initial frame, the frame at t = 2 μs, and the final frame of system 4. The receptor cartoon conformations in each of panels (C-E) are colored by cluster assignments, and the RMSD and residue pair Cα–Cα distance profiles are colored according to their corresponding systems, as shown in panel (A). Arrows in the RMSD and residue pair Cα–Cα distance plots indicate conformational state transitions over the course of the simulation, with the corresponding simulation times marked. Cα atoms are shown as spheres, and side chains are shown as sticks.

System 2, initiated from the G-protein coupled state but without the G-protein, relaxes from the fully active state (cluster a) into cluster i. The SHAP analysis indicates that residue pairs on TM3–TM5, TM3–TM6, and TM4–TM6 discriminate between clusters a and i (**Fig. 4B panel I** and **fig. S12A panel I**), and **Fig. 4C** summarizes the structural changes that accompany the transition from cluster a to i. Residues all along TM3 are identified as significant (**fig. S12A panel I**); analysis of the trajectory shows that this is because TM3 is displaced downward (**Fig. 4C panel I**), which affects all pairwise distances involving residues on TM3. In contrast, the key features involving TM6 are mostly localized to the intracellular end (**fig. S12A panel I** and **fig. S13**), reflecting an inward shift of the intracellular end of TM6 that reduces the distances between the intracellular ends of TM3 and TM6 (**Fig. 4C panel II**) and also between TM4 and TM6 (**Fig. 4C panel III**). Inspection of residues highly ranked by the SHAP analysis reveals that TM6 also undergoes a rotation which changes the conformation of the “P^5.50^I^3.40^F^6.44^” motif (**Fig. 4C panel IV**). All of these structural changes occur simultaneously (see RMSD traces in **Fig. 4C**), suggesting that the conformations in cluster i are strikingly similar to the “pseudo-active” conformation first reported by D’Amore, et al. (*35*).

System 3, initiated from the G-protein coupled state but with neither the G-protein nor the agonist NECA, relaxes from the fully active state (cluster a) into cluster f (**Fig. 4A**). The SHAP analysis indicates that residue pairs on TM1–TM6, TM1–TM7, TM2–TM6, and TM2–TM7 discriminate between clusters a and f (**Fig. 4B panel II** and **fig. S12A panel II**). Residues highly ranked by the SHAP analysis on TM7 localize to the NPxxY motif, which “untwists” at 0.56 μs, reflected by an increase in its RMSD (**Fig. 4D panel I**). A similar structural transition has also been observed in deactivating simulations of β_2_-adrenergic receptor (β_2_AR) (*31*, *36*). In parallel, an increase in interhelical alpha carbon distances involving TM2–TM7 and TM1–TM7 pairs is observed (**Fig. 4D panel II** and **fig. S12B**), associated with the NPxxY transition and an outward displacement of the TM2 kink region, including residues from A50^2.48^ to I60^2.58^ (**Fig. 4D panel III**). A similar transition was reported in previous β₂AR activation/deactivation studies, using the D^2.50^–N^7.49^ distance to identify the transition (*34*). In addition to the TM2 kink, the C^6.47^W^6.48^xP^6.50^ motif (*37*, *38*) undergoes a transition coincident with the others, driving a conformational change in the TM6 kink region (**Fig. 4D panel IV**). These transitions are all consistent with the receptor shifting toward a more inactive conformation.

Changing the membrane environment has a significant effect on the conformational space sampled by the protein. System 4 is initiated from the same structure as system 2, but is embedded in a membrane containing only a zwitterionic lipid (POPC). At 0.69 μs into the simulation the protein switches to a new conformation (cluster g), and subsequently transitions to cluster h at 2.61 μs, both of which are distinct from all of the other states. Residue pairs on TM3–TM7 and TM5–TM7 (**Fig. 4B panel III** and **fig. S12A panel III**) delineate the first conformational transition. As in the transition from the active to pseudo-active state (**Fig. 4C panel I**), TM3 is shifted downward by approximately 1.2 Å (**Fig. 4E panel I**). The downward movement of TM3 increases the distances between TM3 and TM7 residues (**fig. S12C panel I**). TM5–TM7 residue pair distances are also increased, notably V186^5.47^–I274^7.39^ (**Fig. 4E panel II** and **fig. S14**). (Previous studies of β₂AR identified the S^5.46^–G^7.41^ closest heavy atom distance as a key structural feature (*34*).) The TM5 kink region, involving C185^5.46^ and V186^5.47^—one helical turn above P189^5.50^—becomes straighter in cluster g (**Fig. 4E panel II** and **fig. S12C panel II**). Concurrently, the slight outward motion of the TM7 extracellular end also contributes to the increased TM5–TM7 and TM3–TM7 distances.

The second conformational transition from cluster g to cluster h is described by residue pairs on TM2–TM6, TM2–TM7, and TM3–TM6 (**Fig. 4B panel IV** and **fig. S12A panel IV**). Several residues identified by the SHAP analysis localize to the proline kink on TM2 (especially residues from V55^2.53^ to I60^2.58^). The RMSD of the TM2 kink region increases gradually starting at 0.69 μs, followed by a sharp rise at 2.61 μs (**Fig. 4E panel III**), corresponding first to an upward movement of TM2 (**fig. S12D panel I**) that is coupled to the downward shift in TM3, then to a straightening of the TM2 kink. The state switching from cluster g to cluster h is also characterized by an inward bending of the TM6 extracellular end, leading to reduced TM3–TM6 and TM2–TM6 distances, exemplified by the shortened Y103^3.51^–W246^6.48^ (**Fig. 4E panel IV**) and F44^2.42^–L249^6.51^ (**fig. S12D panel II)**. Noting that Y103^3.51^ belongs to the conserved D(E)^3.49^R^3.50^Y^3.51^ motif (*39*, *40*), analysis of the TM3-TM6 ionic lock distance shows that the inward shift of the TM6 intracellular end occurs at 3 μs (**fig. S12D panel III**)—subsequent to the transition event at 2.61 μs. This suggests that the motion of the TM6 extracellular end precedes and perhaps “triggers” the subsequent inward shift of its intracellular end. This observation is consistent with a previous report (*38*), in which upon agonist binding residue C^6.47^ modulates the configuration of the P^6.50^ kink and the subsequent movement of the TM6 cytoplasmic end of β₂AR.

### Arrestin coupled receptor conformations are in the fully active cluster

Beyond the classical paradigm of GPCR signaling, where G-proteins are the primary transducers, certain ligands promote receptor engagement with arrestins over G-proteins. Arrestin binding can sterically hinder G protein coupling, resulting in receptor desensitization (*41*) or internalization (*42*, *43*), and simultaneously trigger distinct arrestin-mediated signals (**Fig. 5A**) (*44–47*). Phosphorylation of the receptor C-terminus by GPCR kinase (GRK) and a helical ICL2 have been identified as key structural determinants for arrestin recruitment in rhodopsin-arrestin complex studies (*4*). Although structural characterization of GPCR–transducer complexes is challenging, several high resolution structures have been published of GPCRs coupled to arrestins (*48–53*), and several authors have used these data to understand arrestin-biased signaling. We therefore asked where in the UMAP-reduced conformational landscape the arrestin bound conformations are found.

**Fig. 5.**
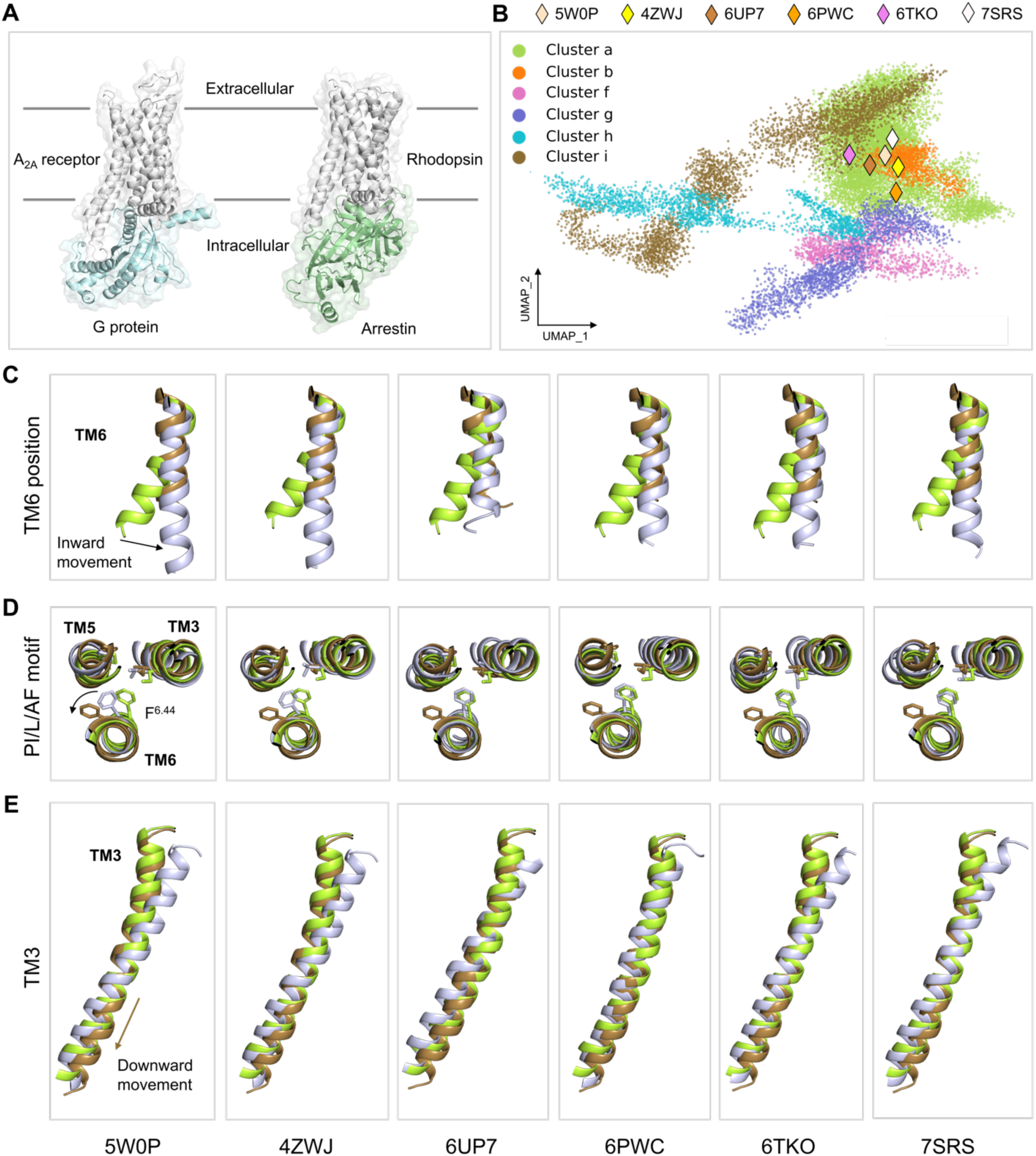
Structural comparison of A_2A_R and arrestin-bound receptors. (**A**) Structural representations of the G-protein bound A_2A_R and arrestin-bound rhodopsin. (**B**) HDBSCAN clustering on the UMAP embedding including six class A arrestin-bound receptors represented by diamonds. Rhodopsin, PDB ID: 5W0P, 4ZWJ; NTSR1, PDB ID: 6UP7, 6PWC; β_1_AR, PDB ID: 6TKO; 5-HT_2B_R, PDB ID: 7SRS. (**C-E**) Structure comparison of each arrestin-bound receptor (lightblue) with the cluster a conformation (initial frame of system 2) and the cluster i conformation (final frame of system 2): (**C**) position of the TM6 intracellular end, (**D**) residue F^6.44^ in the PI/L/AF motif, and (**E**) TM3. Side chains are shown as sticks.

The crystal structure of A_2A_R in complex with arrestin has not yet been determined. To provide a structural reference, we analyzed six arrestin-bound class A GPCR structures: rhodopsin (PDB ID: 5W0P (*48*), 4ZWJ (*49*)), neurotensin receptor 1 (NTSR1) (PDB ID: 6UP7 (*50*), 6PWC (*51*)), β_1_-adrenergic receptor (β_1_AR) (6TKO (*52*)), and 5-hydroxytryptamine (5-HT) serotonin receptor (5-HT_2B_R) (7SRS (*53*)). Based on their inverse Cα–Cα distance features, the six arrestin-bound GPCR structures are all located within cluster a, which also includes the fully active, G-protein coupled conformational ensemble (**Fig. 5B**). (A supplementary movie illustrating the positions of these six structures is available as **Movie S2**, which shows the clusters clearly separated in three UMAP dimensions.) Notably, however, the arrestin coupled structures are all close to the boundary between the fully active conformations (cluster a) and cluster i.

As noted in above, cluster i shares many features with the “pseudo-active” structure described by D’Amore et al. (*35*), who suggest that this is the state that couples to β-arrestin. This prompted us to compare the molecular features of six arrestin-bound GPCRs with the conformations of clusters a and i. As described above, three microswitches undergo conformational changes during the transition from cluster a to cluster i (**Fig. 4C**): an inward movement of the TM6 intracellular end, a counterclockwise rotation of the PIF motif residue F242^6.44^, and a downward shift of TM3. The cytoplasmic end of TM6 in each arrestin-bound GPCR adopts a conformation similar to cluster i (**Fig. 5C**), shifted inward relative to the fully active state cluster a. In the two rhodopsin structures (5W0P and 4ZWJ) the F^6.44^ residue of the PIF motif occupies a position intermediate between clusters a and cluster i, whereas in the remaining four receptors (6UP7, 6PWC, 6TKO, 7SRS), it remains aligned with cluster a (**Fig. 5D**). We observe that the TM3 helix of NTSR1 (6UP7) shows no downward displacement relative to the conformation in cluster a, whereas in NTSR1 (6PWC) the entire TM3 is shifted downward. In contrast, the remaining four receptors exhibit downward movement restricted to the intracellular end of TM3 (**Fig. 5E**). Relative to clusters a and i in this work the arrestin-bound receptors are not exclusively in cluster a (fully active state) or cluster i (pseudo-active state) when assessed via three key microswitches, instead exhibiting molecular characteristics of both states. This is all the more notable, given that the dimensionality reduction and clustering performed in this work does not include any arrestin coupled structures or simulations; with the inclusion of such data the arrestin coupled conformations may well be assigned to their own cluster, perhaps more aligned with cluster i.

## Discussion

With the relentless advance of hardware and software for biomolecular simulation it has become routine to obtain simulation trajectories microseconds in duration, and to launch simulation campaigns that generate hundreds of gigabytes to terabytes of raw data. This presents qualitatively new opportunities to test hypotheses regarding mechanism, by compiling datasets that cover distinct parts of the conformational space of a protein, and then identifying structural motifs that discriminate between functionally significant locales in the conformational landscape. In our view, the discovery of such motifs should be pursued in an investigator-blind way, because experimental design which begins with an expectation that a particular structural element undergoes some conformational change risks confirmation bias; it also might easily miss important but as-yet undescribed structural motifs.

This motivated us to develop an analysis pipeline that takes in molecular simulation data and outputs structural features that discriminate among neighborhoods of similar conformations. In our application to a set of GPCR simulation data we first featurized each sampled protein structure by the inverse alpha-carbon distances between all pairs of residues in the transmembrane bundle. UMAP was used to learn a low dimensional manifold; UMAP has the desirable property that the global structure of the dataset (i.e., which points are close to which other points) is preserved in the lower dimensional space. The data were then clustered with HDBSCAN; inspection of the clusters suggested that they correspond to physically meaningful conformational states. To determine which alpha carbon distances discriminate between clusters a classifier (XGBoost) was trained to discriminate between pairs of clusters, taking as input all residue pair distances. The discriminatory power of each of these distance features was then ranked by an exact interpretability analysis. This constitutes the output of the investigator-blind pipeline: A ranked list of residue pairs that are informative about cluster identity.

This approach was used on a set of simulation data totalling over 28 microseconds of the A_2A_ adenosine receptor, with simulations initiated from different receptor states — fully active, bound to antagonist, etc. Several additional initial states were created by deleting ligands or bound G-protein, with the expectation that these perturbed states would relax into new regions of conformation space. The clusters identified by the UMAP and HDBSCAN steps partition the conformational space in physically sensible ways. For example, two different simulations of antagonist bound receptors are close to one another in the UMAP projection (systems 7 and 9 in **Fig. 1**), while all of the simulations that begin from a fully active structure are initially in the same cluster (cluster a in **Fig. 1**). However, after perturbation (by deleting the agonist and/or G-protein) they all relax into distinct regions of conformation space.

Although a list of alpha carbon distances is not terribly informative on its own, when the top ranked features were mapped onto the structure of the receptor they identified structural changes and motifs (sometimes called “microswitches”) that have previously been established. Comparison of antagonist bound, intermediate, and fully active conformational ensembles with our blind analysis pipeline identified the residues of several microswitches previously reported to be involved in activation: the N^7.49^P^7.50^xxY^7.53^ motif, the R102^3.50^–E228^6.30^ ionic lock, and the D(E)^3.49^R^3.50^Y^3.51^ motif (**Fig. 2**). The analysis also pinpointed changes in interhelical Cα–Cα distances that are known markers of activation, particularly the outward movement of TM6 and the Y^5.58^–Y^7.53^ distance (*54*). Comparison of an intermediate, agonist bound ensemble to one in which the agonist has been deleted identified changes in the extracellular portion of the transmembrane bundle associated with the loss of ligand-protein contacts (**Fig. 3**).

A series of simulations were initiated from the fully active structure, but perturbed by deleting the bound G-protein, bound agonist, and by solvating the receptor in a different membrane environment. Again the blind analysis pipeline localized key features to several known microswitches: the N^7.49^P^7.50^xxY^7.53^ motif, the P^5.50^I^3.40^F^6.44^ motif, the D(E)^3.49^R^3.50^Y^3.51^ motif, and the C^6.47^W^6.48^xP^6.50^ motif (**Fig. 4**) (*25*, *34–36*, *38*). Of particular note is the relaxation of the receptor when the G-protein is deleted (but NECA remains bound) into a “pseudo-active” conformation previously reported by D’Amore, et al. by a rotation of the PIF motif (*35*). This state appears to be similar to arrestin bound structures of other GPCRs (**Fig. 5**), which our clustering analysis places on the boundary of the pseudo-active cluster.

The real value of an investigator blind analysis is revealed by the discovery of new structural features associated with transitions among different conformational ensembles. In several instances our analysis points to a kink in TM2 near the conserved residue D^2.50^, which undergoes a straightening as the receptor relaxes from a fully active or active intermediate conformation after deletion of agonists (**Fig. 3C**, **Fig. 4D**, **Fig. 4E**). Residue D^2.50^ binds a sodium ion to stabilize the inactive state (*55*), has been reported by NMR to undergo a conformational change during ligand binding to A_2A_R (*56*, *57*), and has been reported to be a key titratable residue for β_2_AR activation (*34*, *36*). Our analysis pipeline also identifies coupled motions of TM2 and TM3, in which TM2 moves up as TM3 moves down upon inactivation, supporting a prior report by D’Amore et al (*35*).

This work considers a set of simulation data for a single receptor totalling a few tens of microseconds. What new features might be discovered by applying a similar approach to a far larger dataset encompassing many different GPCRs, as are available in the GPCRmd database (*6*)? We anticipate that this will require a different, much less computationally demanding approach for dimensionality reduction and clustering, perhaps leveraging recent advances for rapid comparison of protein structures that scale to hundreds of millions of structures (*58*).

Beyond the structure-function details of interest to the GPCR community, we hope that more simulation groups take up an “experimental” design like the one put forth here, which we believe may reduce confirmation bias. Instead of beginning the analysis of the data from an *expectation* that a particular motif changes its conformation, the data (and its blinded analysis) show the investigator where to look.

## Materials and Methods

### System setup for molecular dynamics simulations

Simulations were initiated from the coordinates of four different crystal structures of the adenosine A_2A_ receptor (A_2A_R): the receptor in complex with the agonist NECA and an intracellular engineered G protein (PDB: 5G53) (*59*), the receptor in complex with a triazole-carboximidamide (TC) antagonist (PDB: 5UIG) (*60*), the receptor in complex with CGS21680 (PDB: 4UHR) (*61*), and the receptor in complex with ZM241385 (PDB: 3PWH) (*62*). We created two additional starting configurations from 5G53: one in which the G-protein was deleted, and one in which both the G-protein and NECA were deleted. Apo states were also created from 5UIG and 4UHR. The set of simulated receptors are listed in **Table 1**. Modeller 10.5 (*63*) was used to rebuild residues 147 to 158 and residues 212 to 223 that were not resolved in the receptor PDB structure 5G53, residues 155 to 157 and 263 in the PDB structure 4UHR, residues 149 to 158 and 209 to 218 in the PDB structure 5UIG, and residues 150 to 157 in the PDB structure 3PWH. The protein was embedded in an asymmetric membrane model (MM) (*64*), in which the membrane composition contains 12 lipids and is a mixture of sphingolipids (SM), phosphatidylcholine (PC), phosphatidylethanolamine (PE), ether-linked (plasmalogen) PE (abbreviated PLAS), phosphatidylserine (PS), and cholesterol. The asymmetric membrane lipid compositions are listed in **Table S9**. One additional system was built in which A_2A_R in complex with NECA was embedded in a POPC bilayer.

Each system was prepared using the membrane builder in CHARMM-GUI (*65*). The protein complex was embedded in the membrane lipids mentioned above, solvated with TIP3P (*66*) water molecules to provide a 30 Å-thick layer above and below the bilayer. Na^+^ ions were added to neutralize the system, and additional Na^+^ and Cl^−^ ions were added to maintain 0.15 M ionic concentration. The systems with the asymmetric MM contained approximately 430,000–500,000 atoms and measured around 18.5 × 18.5 × 15.3 nm^3^. The system with the POPC bilayer contained 112,873 atoms and measured 9.5 × 9.5 × 14.0 nm^3^. Protein and lipids were modeled with the CHARMM36 force field (*67*, *68*), and the ligands were modeled with the CHARMM general force field (*69*).

### Simulation details

The equilibration of the system 6 in **Table 1** was run with NAMD version 3 (*70*), following CHARMM-GUI’s default six-step protocol. The initial configuration was relaxed by 10000 steps of steepest descent. The first three simulations had 1 fs timestep for a total of 375 ps (125 ps × 3) of simulation time, and the last three steps had a 2 fs timestep for a total of 1500 ps (500 ps × 3) simulation length. During the six-step equilibration protocol, velocities were reassigned every 500 steps. The heavy atoms of the receptor backbone and the ligand NECA were positionally restrained but with a decreasing force constant from 10.0 to 0.1 kcal mol^−1^ Å^−2^ following CHARMM-GUI’s default. The force constant was gradually reduced from 5.0 to 0 kcal mol^−1^ Å^−2^ for the heavy atoms of the receptor sidechain. Similarly, the harmonic restraints on the heavy atoms of lipid headgroups were reduced from 5.0 to 0 kcal mol^−1^ Å^−2^ in a stepwise manner. The simulation box volume was allowed to change semi-isotropically via a Langevin piston with a barostat damping time scale of 25 fs and an oscillation period of 50 fs (*71*). The target pressure and temperature were 1.01325 bar and 310 K, respectively. All covalently bonded hydrogens were constrained by SHAKE (*72*). A cutoff of 1.2 nm was used for the electrostatic and van der Waals interactions, while the switching algorithm was on if the distance between two atoms was between 1.0 nm and 1.2 nm, ensuring the potential was smoothly reduced to 0 at the cutoff distance. The long-range electrostatic interactions were computed using the Particle Mesh Ewald (PME) method (*73*, *74*) on a 1 Å grid, with a tolerance of 10^−6^ and sixth order interpolation. The nonbonded interactions were not computed only if two atoms were connected within 3 covalent bonds. The neighbor list was updated every 10 steps, including all pairs of atoms whose distances were 1.2 nm or less. The sixth equilibration step was run for another 30 ns to ensure the receptor was well equilibrated. An additional 70 ns production run was performed in which all restraints were removed.

The equilibrated binary restart files from NAMD were converted into DMS format for further production simulation on Anton2. Viparr 4.7.49c7 was used to add force field information. Integration was performed under constant pressure (1 atm), temperature (310 K), and particle number with the multigrator (*75*) method with a 2.5-fs time step. Temperature was controlled by a Nose-Hoover (*76*) thermostat coupled every 24 timesteps and pressure was controlled by the Martyna-Tobias-Klein barostat (semi-isotropic) coupled every 480 timesteps (*77*). Nonbonded interactions were cut off at 1.2 nm. Long-range electrostatic interactions were computed using the u-series method (*78*) following Anton2’s default. Hydrogens were constrained by M-SHAKE (*79*). The production simulation was performed for 5 µsec, and configurations were stored every 480 ps.

The other systems in **Table 1** were run with GROMACS version 2024 (*80*, *81*). These systems were subjected to a series of energy minimization and equilibration steps with the input files generated from the CHARMM-GUI membrane builder (*65*). The CHARMM36 force field parameters were used for protein, lipids, salt (0.15 M NaCl), and explicit TIP3P water (*68*). The ligand NECA, CGS21680, TC, and ZM241385 was modeled with the CHARMM general force field (*69*). The initial configuration was relaxed by 10000 steps of steepest descent. The first two simulations had 1 fs timestep for a total of 500 ps (250 ps × 2) of simulation time in isothermal−isochoric (NVT), the third step had 1 fs timestep for 250 ps in semi-isotropic, and the last three steps had a 2 fs timestep for a total of 1500 ps (500 ps × 3) simulation length in semi-isotropic. The heavy atoms of the receptor and ligand, and the phosphorus atom of lipids were positionally restrained but with a decreasing force constant in a stepwise manner following CHARMM-GUI’s default. All restraints were removed during the production MD. The temperature was maintained at 310 K using the v-rescale thermostat (*82*) with τ_t_ = 1.0 ps. In the preproduction and production semi-isotropic run, the pressure of 1 bar was maintained using C-rescale (*83*) with τ_p_ = 5.0 ps and compressibility of 4.5 × 10^−5^ bar^−1^. The production ran for 2 µs for systems 1, 3, and 8, 4.4 µs for the system 4, and 1 µs for systems 5, 7, and 9. Orthorhombic periodic boundary conditions were applied to each system. The nonbonded interaction neighbor list was updated every 20 steps. A 1.2-nm cutoff was used for the electrostatic and van der Waals interactions. The long-range electrostatic interactions were calculated by using the Particle-Mesh Ewald method (*73*, *74*) after a 1.2-nm cutoff. The bonds involving hydrogen atoms were constrained using the linear constraint solver (LINCS) algorithm (*84*).

### Data preprocessing and simulation analysis

For GPCR conformations along the simulation trajectories, we isolated the seven transmembrane alpha-helices, excluding both intracellular and extracellular loops. The residues involved in the feature calculations are listed in **Table S10**. We then calculated the inverse Cα-Cα atom distances between helices, resulting in a 28,400 x 15,916 matrix, where 28,400 corresponds to the number of conformations (one snapshot per nanosecond) across nine simulations, and 15,916 represents the number of alpha-carbon pairwise distances.

PyMOL (*85*) and VMD (Visual Molecular Dynamics) (*86*) were used for visualizing the MD trajectories and image rendering. Data analysis and plotting were performed using in-house python scripts based on publicly hosted python packages: NumPy (*87*), MDAnalysis (*88*), and matplotlib (*89*).

### Dimensionality reduction and clustering

Dimensionality reduction aims to transform data into a relevant and lower-dimensional space by uncovering its intrinsic structure and minimizing redundancy. This process helps retain informative features while filtering out noise and correlated features. We derived the conformations from MD simulations of A_2A_ receptors. To project the conformations onto a lower-dimensional manifold, we used three different dimensionality reduction methods, principal component analysis (PCA) (*90*), t-distributed stochastic neighbor embedding (t-SNE) (*16*), and uniform manifold approximation and projection (UMAP) (*20*, *21*). The calculated 15,916 inverse Cα-Cα atom distances were used as input features for dimensionality reduction. PCA was available at https://scikit-learn.org/stable/ with parameter settings as n_components=100, svd_solver=‘auto’, iterated_power=‘auto’, random_state=None. T-SNE was available at https://scikit-learn.org/stable/ with parameter settings as n_components=2, perplexity=30, init=‘random’, max_iter=1000, metric=’’euclidean”, random_state=0.

UMAP was available at https://github.com/lmcinnes/umap/. UMAP first constructs a high dimensional weighted graph representation of the data where the edge weights represent the likelihood that pairs of points are connected. A cost function is then applied to optimize a low-dimensional representation that maintains the original structural similarity. Two key input parameters, the number of nearest neighbours and the minimum distance, control the trade-off between local and global structures. The number of nearest neighbours defines how many nearest neighbors are considered to obtain the initial high-dimensional graph, and the minimum distance sets the minimum distance between points, controlling how tightly points are packed in the resulting low-dimensional embedding representation. For UMAP, hyperparameter space (n_neighbors: [50, 100, 200, 300, 400, 500], n_components: [2, 3, 4, 5, 6, 7, 8, 9, 10], and min_dist: [1.0]) was tested to find the optimal hyperparameter via a grid search. Here, the min_dist value was set to 1.0 to prevent UMAP from packing conformations together and would focus on the preservation of the broad topological structure. The resulting embedding was subjected to hierarchical density-based spatial clustering of application with noise (HDBSCAN) (*91*) clustering to identify distinct conformational ensembles. For HDBSCAN, a grid search over min_cluster_size of [100, 300, 500, 700, 1000] and min_smaples of [10, 50, 100] were tested with approx_min_span_tree=False and gen_min_span_tree=True. The Density Based Cluster Validity (DBCV) score (*92*) was used to evaluate the hyperparameter choices for UMAP and HDBSCAN clustering (**Table S1**). A variety of unsupervised clustering metrics, Silhouette Coefficient (SC) score (*93*), Calinski-Harabasz (CH) index (*94*), and Davies-Bouldin (DB) score (*95*), are available in scikit-learn (https://scikit-learn.org/stable/). Their drawbacks are generally higher for convex clusters such as density-based clusters like DBSCAN or HDBSCAN as scikit-learn mentioned, and thus they are not used in this study.

### Classification and feature importance interpretation

Multiple conformational states (e.g., active, intermediates, inactive) of A_2A_ receptors exist throughout simulations, it is important to understand which input features most strongly distinguish these states. Research in data science has advanced techniques to compute feature importance. Lloyd Shapley derived a mathematical formula known as Shapley values, rooted in collaborative game theory (*96*). This concept provides a fair distribution of a payout among players working toward a common goal, even when contributions are not equal among players. Strumbelj and Kononenko first introduced the use of Shapley values for ML interpretability in 2010 (*97*), and Lundberg et al. popularized and extended it to the so-called SHAP (SHapley Additive exPlanations) analysis (*30*). SHAP values provide a fair and unbiased measure of each feature’s contribution to the predicted value of each sample. For example, supposing a ML model is trained with the following input data: X_k_={x_1_, x_2_,…, x_n_}^S^. To evaluate the contribution of each input feature to the ML model, the SHAP method uses an explanatory model (F). These are described in the below equations:

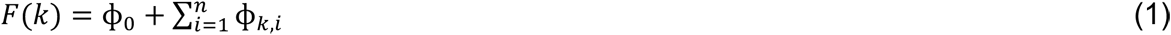

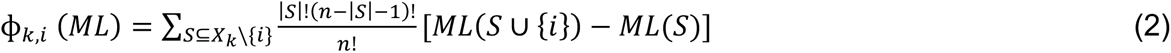

Where ϕ_0_ is the average model output (the baseline), n is the number of input features, ϕ_k,i_ (ML) is the SHAP value (contribution) of the feature i in sample k to the ML model, S is a subset of the feature set that does not include feature i, |S| is the size of subset S, ML(S∪{i}) is the model’s output when adding feature i to subset S, ML(S) is the model’s output using only features in subset S, and \ is the difference set notation for set operations.

Based on the equation definitions, we see for a given sample k, the SHAP values assigned to each feature describe the extent to which that feature contributed to the difference between the individual prediction of this sample from the average model prediction. It is often computationally intensive using the equations mentioned above to calculate SHAP values for feature contributions of a ML model prediction. To address this, various implementation and approximation techniques have been developed to make SHAP value calculation more efficient. For neural network-based models, these approximations typically rely on the model differentiability or the computational graph (*98*). In contrast, tree-based models—such as XGBoost, random forest, CatBoost—allow for the exact computation of SHAP values by leveraging their tree structure (*99*).

XGBoost, which stands for eXtreme Gradient Boosting, is an evolving supervised machine learning (ML) algorithm developed by Chen and Guestrin and the XGBoost algorithm is thoroughly described in their publication (*29*). XGBoost was available at https://xgboost.readthedocs.io/en/release_3.0.0/. To capture key feature changes associated with A_2A_R conformational state transitions, we performed binary classification on selected pairs of clusters determined by HDBSCAN clustering. The XGBoost classifier was used with parameter settings as n_estimators=2000, max_depth=6, objective=’binary:logistic’, learning_rate=0.01. Using the inverse Cα-Cα atom distances as input and the cluster ID as output, XGBoost classifier was trained. We ran five-fold cross-validation using stratified splits for training/validation set by the ratio of 4:1. We implemented accuracy, precision, recall, F1 score, and confusion matrix as metrics to evaluate the classification performance. SHAP analysis was performed to evaluate XGBoost classifier model and SHAP values were calculated to explain feature importance in each fold.

## Acknowledgements

This research was supported by the US National Institute of General Medical Sciences by award number R35-GM153273. This work used Delta at the National Center for Supercomputing Applications through allocation BIO240227 from the Advanced Cyberinfrastructure Coordination Ecosystem: Services & Support (ACCESS) program, which is supported by U.S. National Science Foundation grants #2138259, #2138286, #2138307, #2137603, and #2138296. Anton2 computer time was provided by the Pittsburgh Supercomputing Center through grant R01GM116961 from the National Institutes of Health. The Anton2 machine was made available by D. E. Shaw Research.

## Author contributions

Conceptualization: J.J. and E.L. Methodology: J.J. and E.L. Investigation: J.J. Data analysis: J.J. Visualization: J.J. and E.L. Supervision: E.L. Project administration: E.L. Writing—original draft: J.J. and E.L. Writing—review and editing: E.L.

## Data, code, and materials availability

All data are available in the main text and/or the Supplementary Materials. All raw simulation parameters and trajectories have been submitted to the GPCRmd platform https://www.gpcrmd.org/dynadb/publications/2507/. Please use the following **Dynamics IDs** for systems 1-9: 3231, 3244, 3232, 3233, 3234, 3242, 3236, 3237, 3238, along with the **Submission password**: a2ar_simulation. The data and code used for dimensionality reduction, clustering, classification, and feature importance explanation are available on https://doi.org/10.5281/zenodo.19002785.

## Supplementary Materials

**This PDF file includes**

Supplementary Text

Figs. S1 to S14

Tables S1 to S10

Legends for movies S1 and S2

**Other Supplementary Materials for this manuscript include the following:**

Movies S1 and S2

