## Supplemental materials for "Investigator-blind discovery of structural elements controlling GPCR function"

Jingjing Ji and Edward Lyman*

**This PDF file includes:**

Supplementary Text
Figs. S1 to S14
Tables S1 to S10
Legends for movies S1 and S2

**Other Supplementary Materials for this manuscript include the following:**

Movies S1 and S2

### Supplementary Text

Dimensionality reduction and clustering

PCA has been used for more than 20 years in the simulation field to project high dimensional configuration spaces into lower dimensional representations. When applied to our dataset we found that the first three components only capture about 50% of the observed variance, and twenty dimensions are required to reach 80% of the variance (**Fig. S1**). Seeking a still lower dimensional description we turned to nonlinear methods.

T-SNE is a visualization technique that transforms high dimensional data into a low dimensional embedding, aiming to place similar points close together and dissimilar points far apart. It converts similarities between data points to joint probabilities and is designed to minimize the Kullback-Leibler divergence between the joint probabilities of the low dimensional embedding and the high dimensional data. Local neighborhoods are usually preserved very well, as we see that each of the nine simulation trajectories tends to be broken into multiple separate islands in the t-SNE plot (**Fig. S2**). However, the global structure is often distorted in t-SNE embeddings. For example, conformational ensembles may be artificially pushed apart even if they were close in the high dimensional space, and thus inter-ensemble distances are not meaningful. Because t-SNE does not maintain global structural relationships we reasoned that it may not be well-suited to analyzing a large data set like ours with an eye toward rationalizing structure-function.

UMAP is a manifold learning technique based on Riemannian geometry and algebraic topology. Beginning from the assumption that there exists an underlying manifold structure of the data (a good assumption for a classical system of interacting particles), UMAP is designed to preserve the local and global structure of the original data. In contrast to t-SNE, UMAP tries to maintain relative distances and connectivity between points in the high dimensional manifold when projected into the lower dimensional representation, which means that clusters that are close in high dimensional space remain close in low dimensional embedding. We view these characteristics as desirable for the present application.

Like many other dimensionality reduction algorithms, the UMAP algorithm starts by building a k nearest neighbor graph. The number of nearest neighbors to consider when constructing the graph is a key hyperparameter; it sets the scale over which the manifold is assumed to be locally flat, and thus controls averaging over small scale variations in the hypersurface. Variations below this scale will not be detected, and larger-scale features will be learned by patching together pieces of this scale. A second important hyperparameter is the minimum distance between points after projecting into the reduced dimensional space. A smaller value will yield tight clusters of points after projection, a larger value will distribute them more uniformly and better preserve long range topological structure. The final hyperparameter for UMAP is the dimensionality of the projection.

To perform clustering on the projected conformational space we used hierarchical density-based spatial clustering of applications with noise (HDBSCAN). HDBSCAN is robust for data sets with varying density and for clusters with different shapes, and doesn't require a predetermined number of clusters. HDBSCAN also requires choosing some hyperparameters, the most critical one being the minimum size of a cluster, with clusters below this threshold being treated as noise. The result of a run (i.e., a particular combination of hyperparameters) was evaluated by calculating the density based cluster validity (DBCV), which scores a clustering by comparing within-cluster distances to between-cluster distances. After a thorough search of the hyperparameter space (we considered UMAP and HDBSCAN hyperparameters together during the search) we settled on n_neighbors = 500, n_components = 9, min_dist = 1.0 for the UMAP hyperparameters and min_cluster_size = 1000, min_samples = 10 for the HDBSCAN clustering step (**Table S1**). We then slightly decreased min_dist from 1.0 to 0.95 and observed that ten clusters consistently emerged in the conformational space for min_dist values between 0.99 and 0.96. The final UMAP embedding was obtained using min_dist = 0.99, followed by clustering. Results based on these choices are presented below.

A

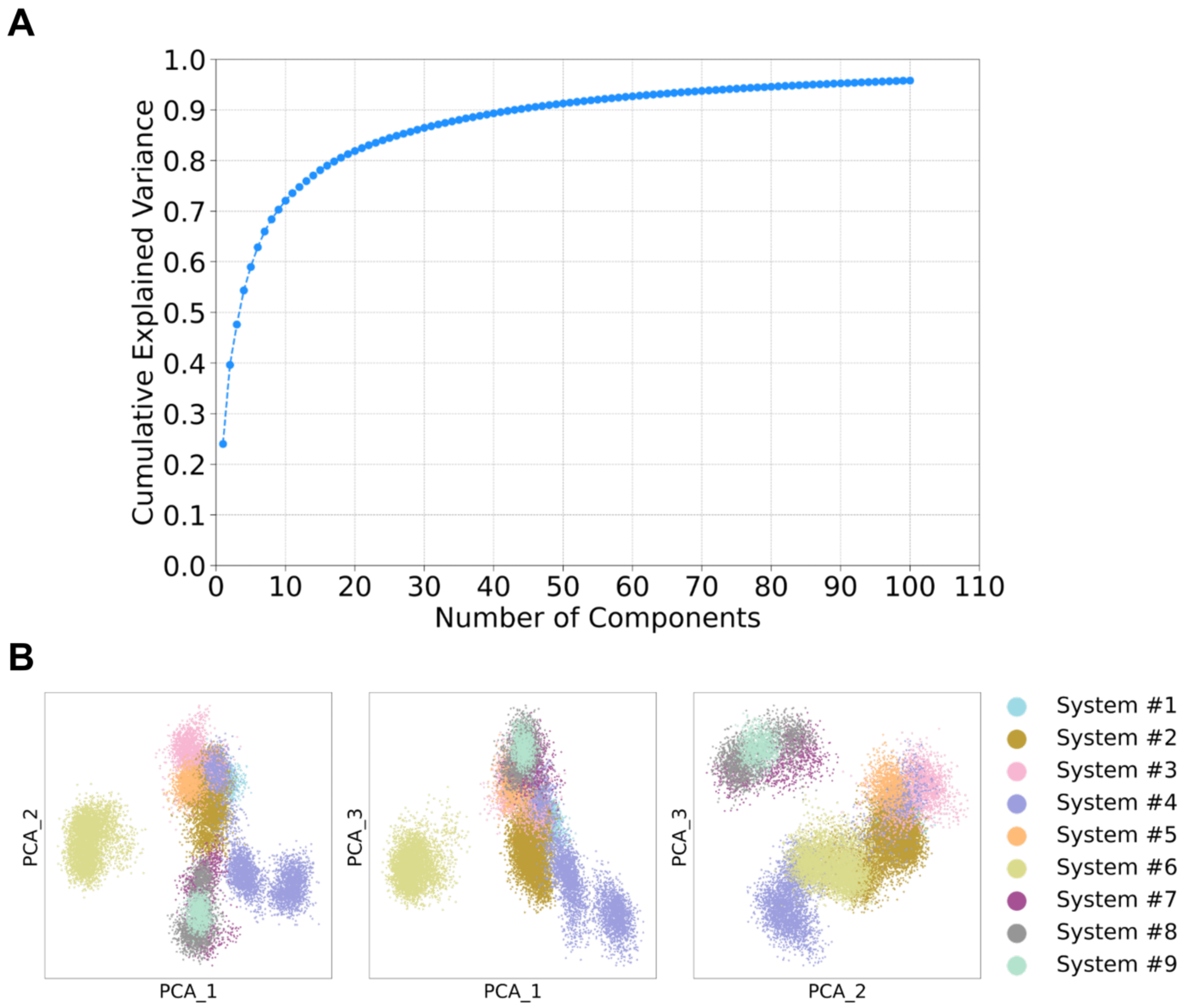


**Fig. S1. PCA analysis across nine simulation trajectories.** (**A**) Cumulative explained variance with the first 100 components. (**B**) PCA projection onto the first three principal components across nine simulation trajectories. Each point represents a simulation snapshot.

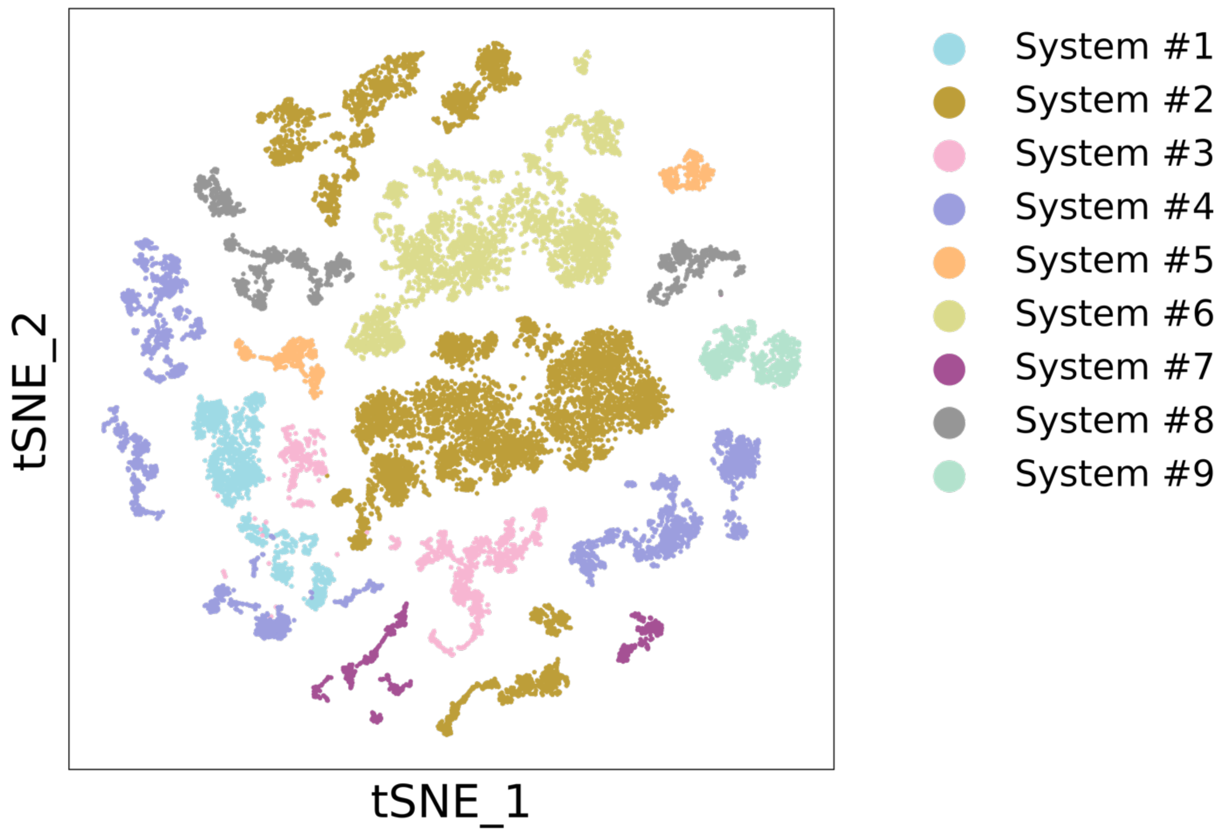


**Fig. S2. T-SNE projection across nine simulation trajectories.** Each point represents a simulation snapshot.

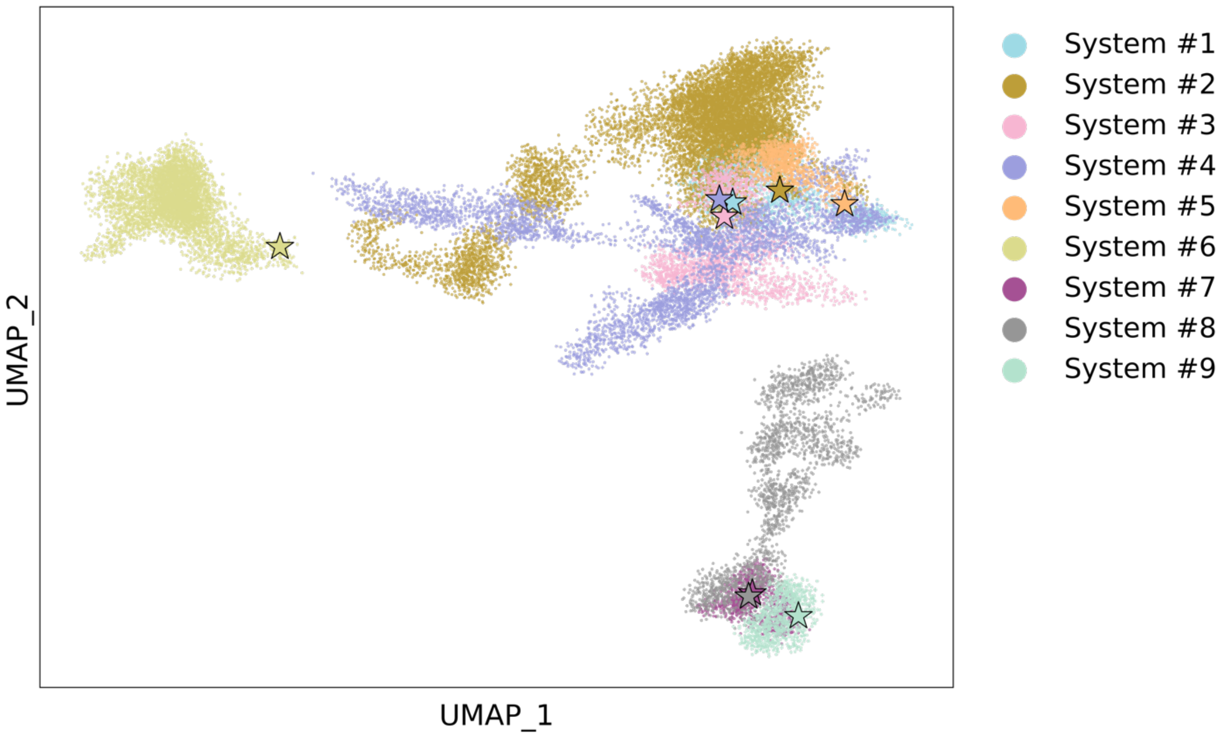


**Fig. S3. UMAP projection across nine simulation trajectories.** Each point represents a simulation snapshot. The stars denote the initial conformations of the nine systems used in the production runs.

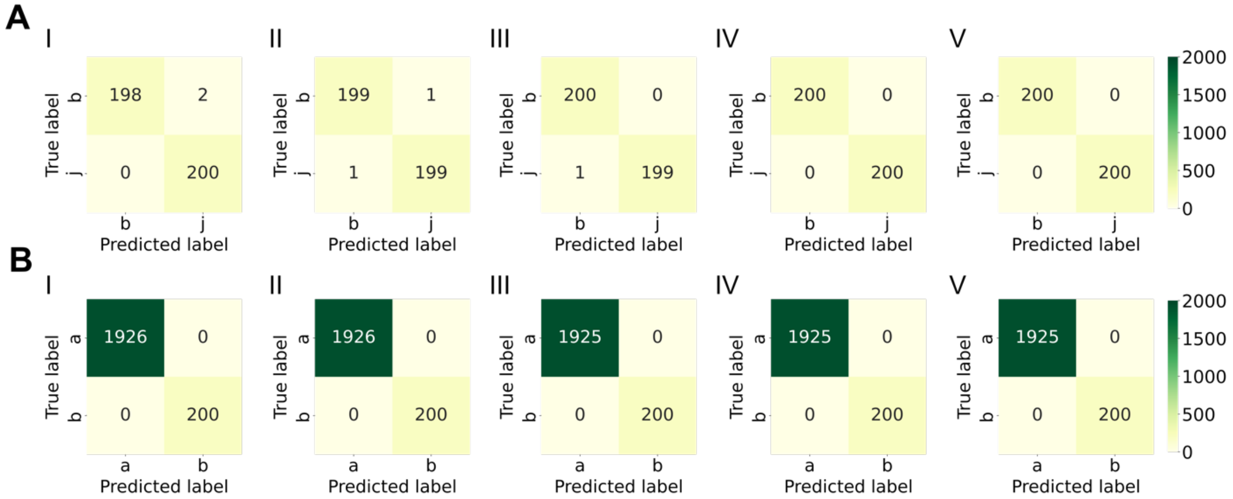


**Fig. S4. Confusion matrices evaluating classification across five cross-validation folds (I-V).** (**A**) clusters j/b, (**B**) clusters b/a.

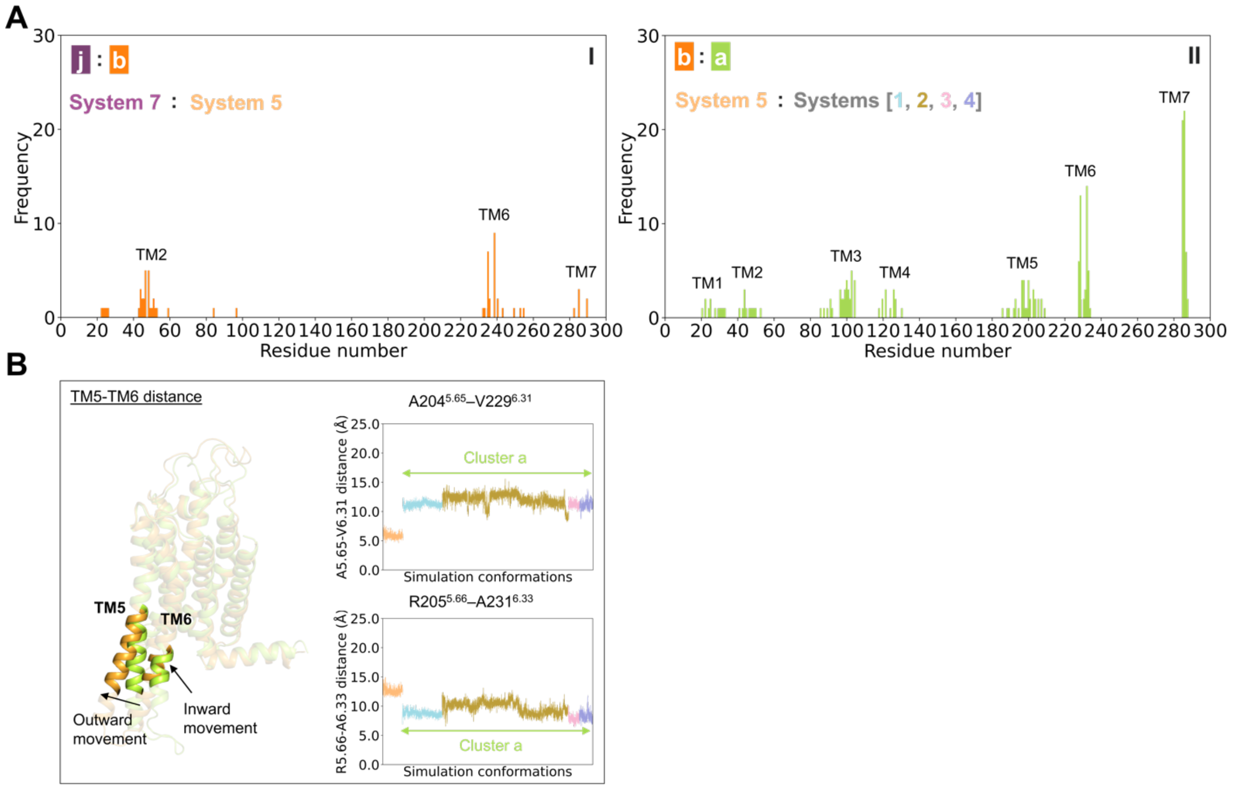


**Fig. S5. Supplementary conformational differences between clusters.** (**A**) Frequencies of residues derived from the top interhelical Cα–Cα pairs identified by SHAP analysis in distinguishing between the two clusters. Panel A-I reflects the top 32 interhelical Cα–Cα pairs, whereas panel A-II reflects the top 100. (**B**) Supplementary features in the insets distinguish clusters a and b. Overlaid receptor conformations correspond to the final frames of systems 1 and 5. The receptor cartoon conformations are colored by cluster assignments, and the residue pair Cα–Cα distance profiles are colored according to their corresponding systems, as shown in panel (A).

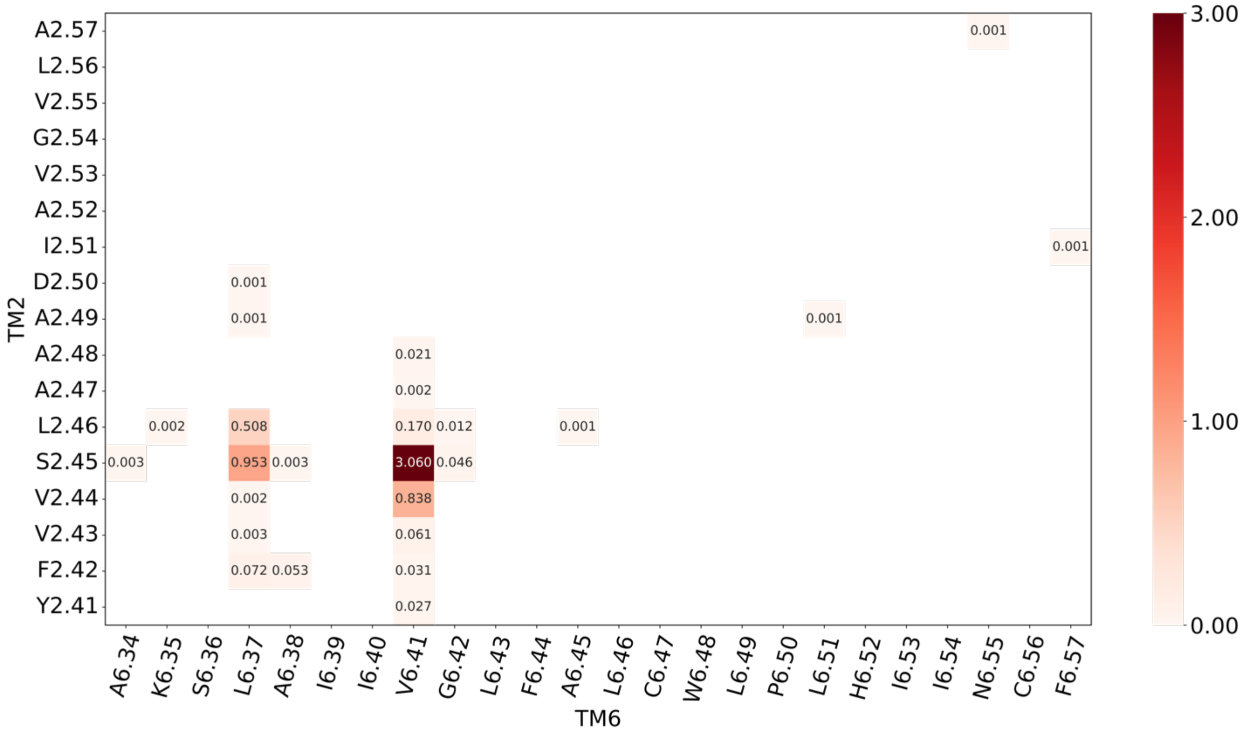


**Fig. S6. SHAP value heat map of residue pairs (Cα–Cα distances) indicating their contributions to distinguish clusters b and j.** Values are the average absolute SHAP values across five cross-validation folds.

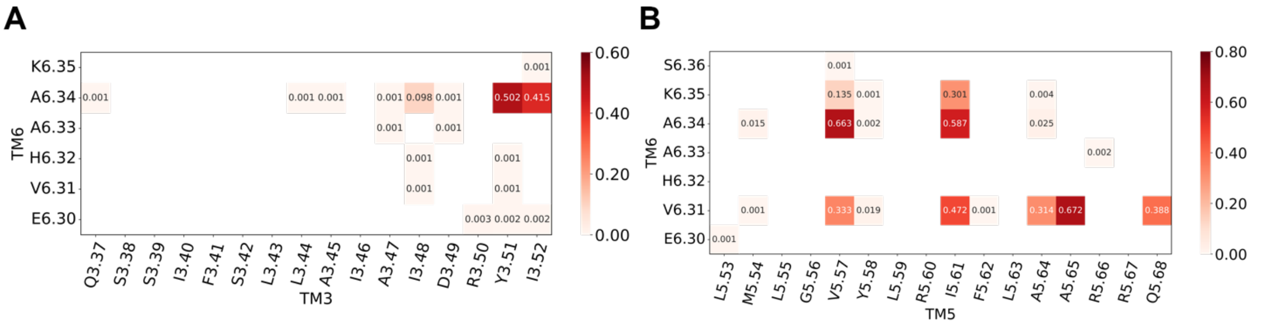


**Fig. S7. SHAP value heat map of residue pairs (Cα–Cα distances) indicating their contributions to distinguish clusters a and b.** Values are the average absolute SHAP values across five cross-validation folds.

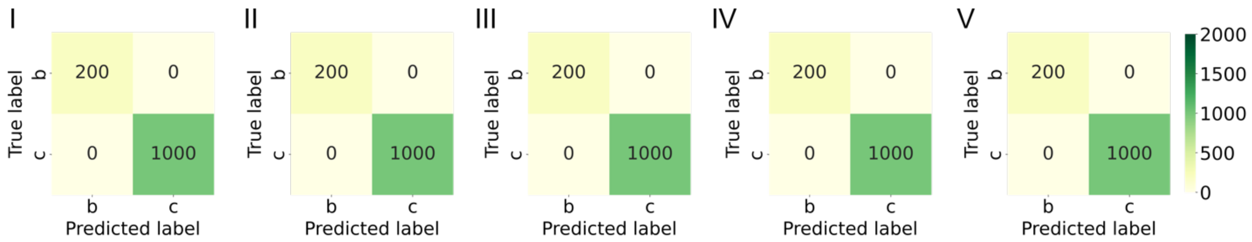


**Fig. S8. Confusion matrices evaluating clusters b/c classification across five cross-validation folds (I-V).**

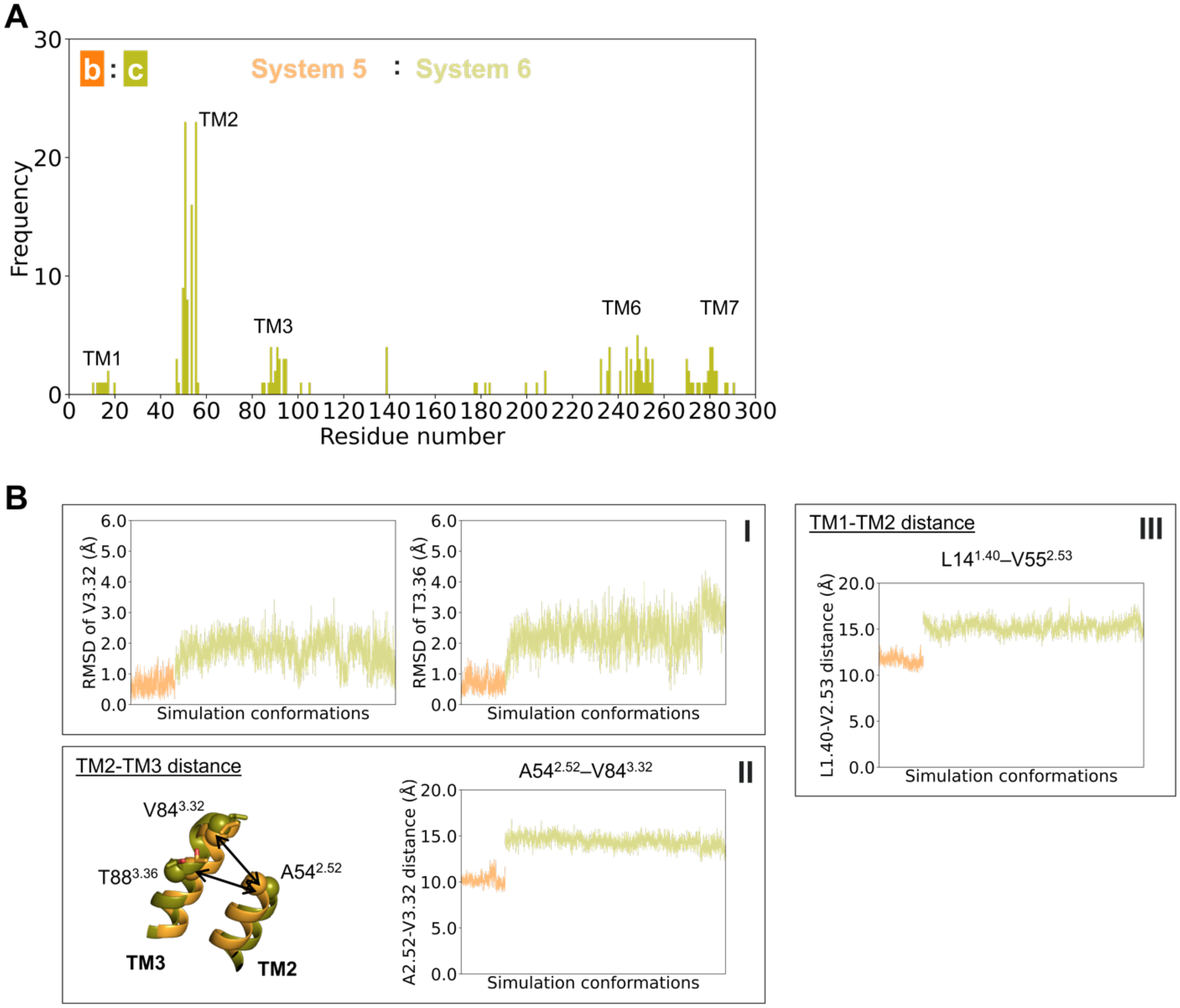


**Fig. S9. Supplementary conformational differences between clusters.** (**A**) Frequencies of residues derived from the top 100 interhelical Cα–Cα pairs identified by SHAP analysis in distinguishing between the two clusters. (**B**) Supplementary features in the insets distinguish between clusters b and c. Overlaid receptor conformations correspond to the final frames of systems 5 and 6. The receptor cartoon conformations are colored by cluster assignments, and the RMSD and residue pair Cα–Cα distance profiles are colored according to their corresponding systems, as shown in panel (A). Cα atoms are shown as spheres.

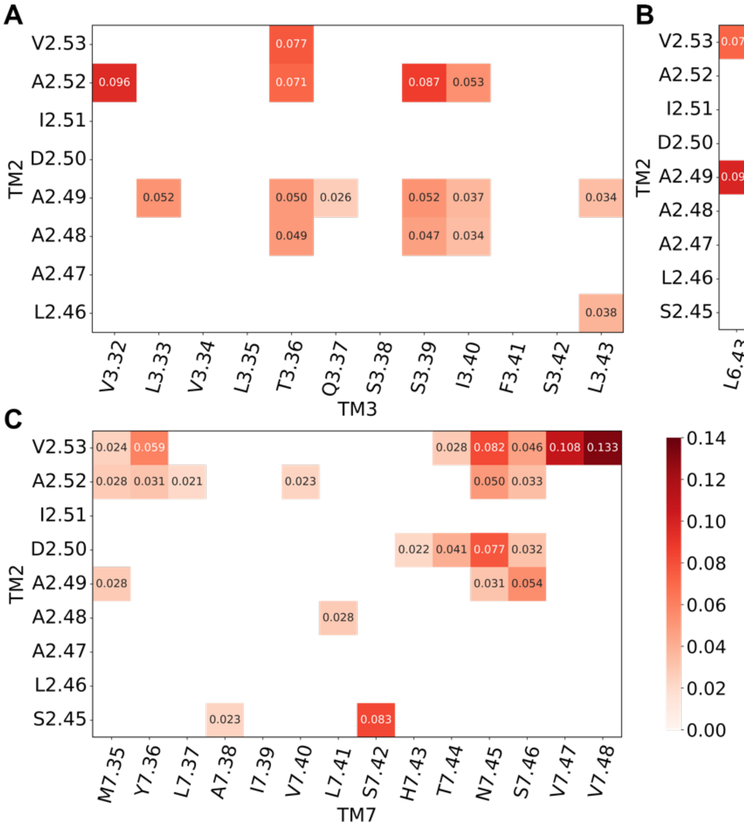


**Fig. S10. SHAP value heat map of residue pairs (Cα–Cα distances) indicating their contributions to distinguish clusters b and c.** Values are the average absolute SHAP values across five cross-validation folds.

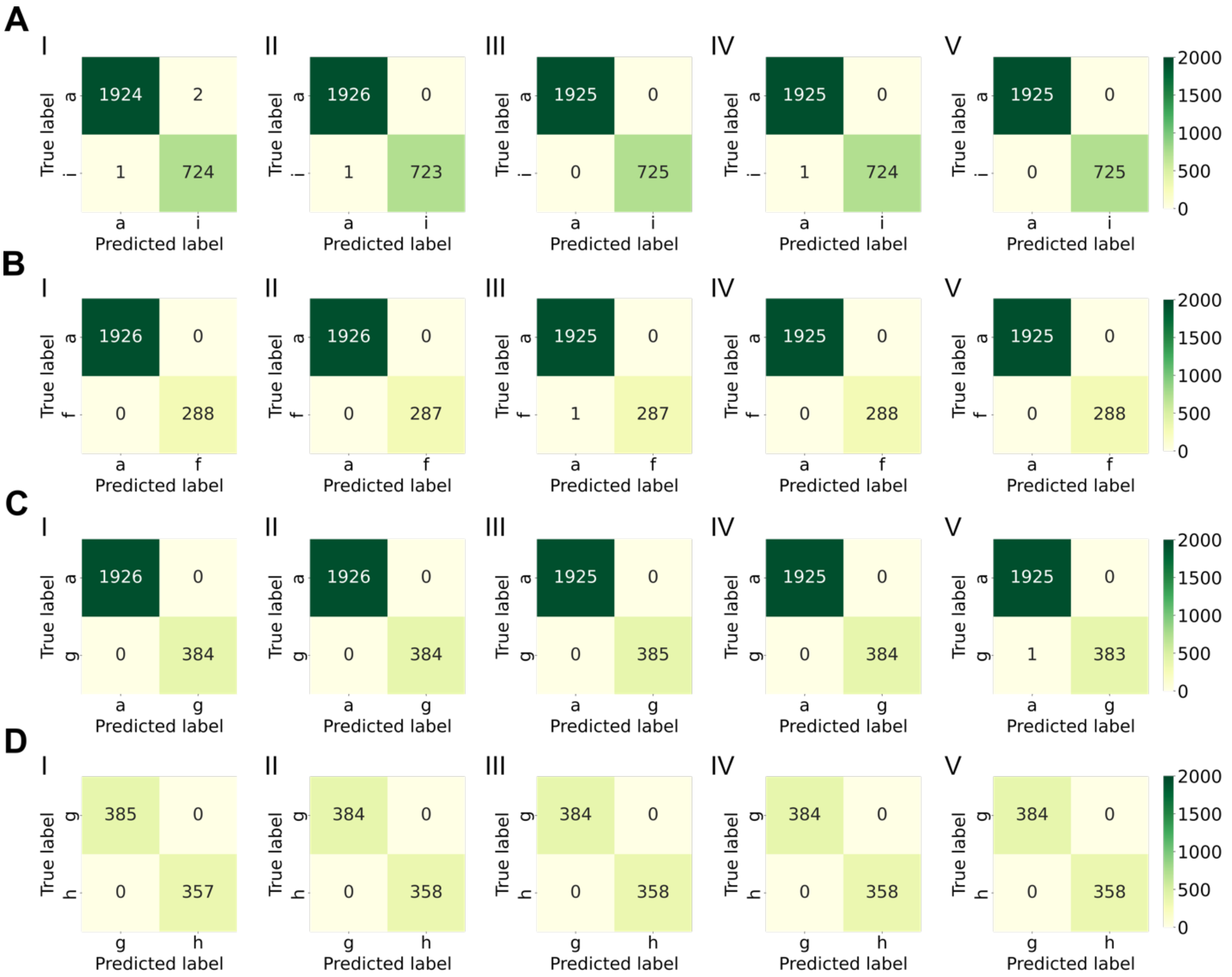


**Fig. S11. Confusion matrices evaluating classification across five cross-validation folds (I-V).** (**A**) clusters a/i, (**B**) clusters a/f, (**C**) clusters a/g, and (**D**) clusters g/h.

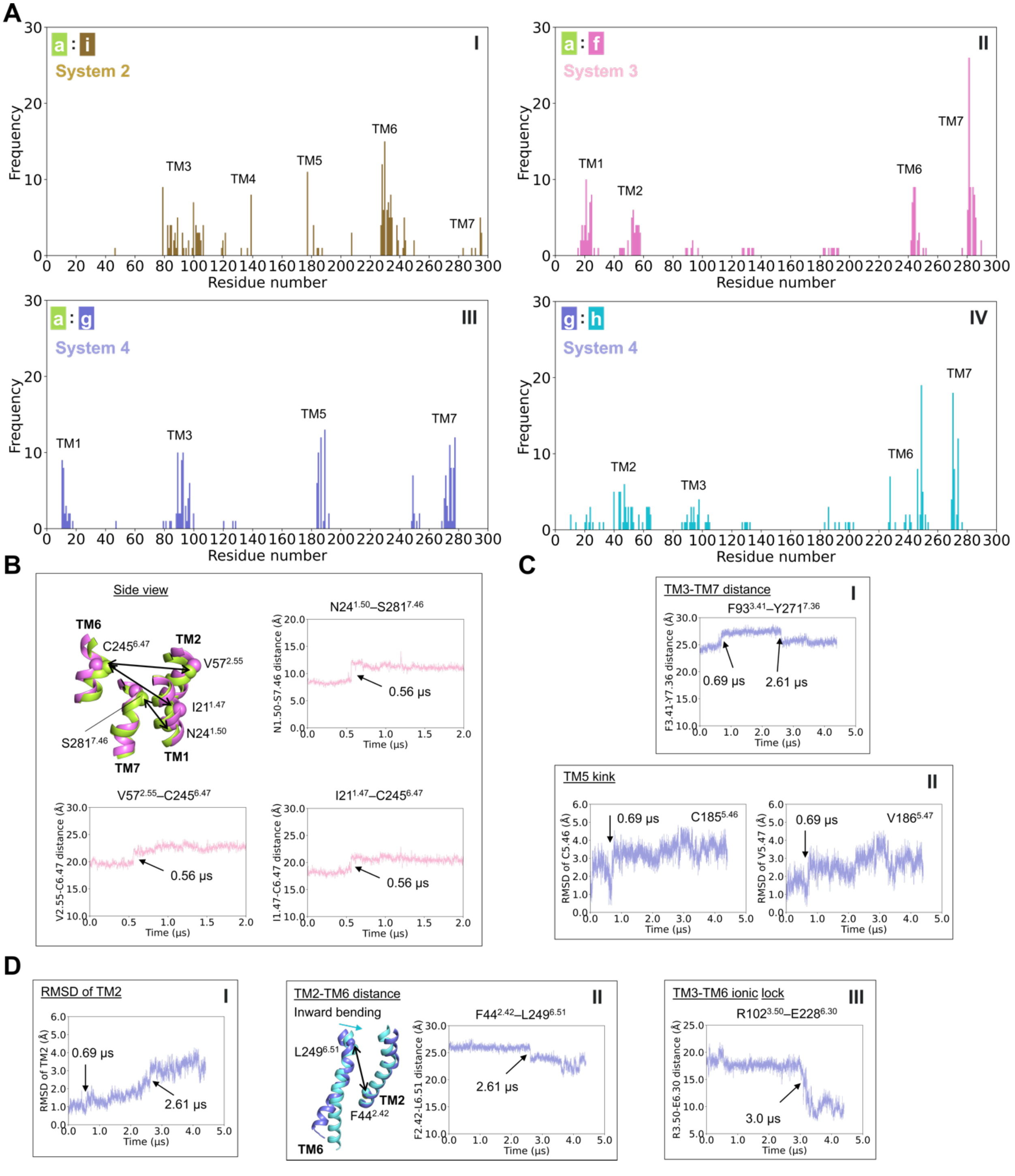


**Fig. S12. Supplementary conformational changes during simulated state transitions.** (**A**) Frequencies of residues derived from the top 100 interhelical Cα–Cα pairs identified by SHAP analysis in distinguishing between the two clusters. (**B-D**) Supplementary feature changes in the insets associated with state transitions in simulations: (**B**) transition from cluster a to f in system 3, and (**C-D**) sequential transitions from cluster a to g and g to h in system 4. Overlaid receptor conformations in (B) correspond to the initial and final frames of system 3; and in (C-D), the initial frame, the frame at t = 2 μs, and the final frame of system 4. The receptor cartoon conformations in each of panels are colored by cluster assignments, and the RMSD and residue pair Cα–Cα distance profiles are colored according to their corresponding systems, as shown in panel (A). Arrows in the RMSD and residue pair Cα–Cα distance plots indicate conformational state transitions over the course of the simulation, with the corresponding simulation times marked. Cα atoms are shown as spheres.

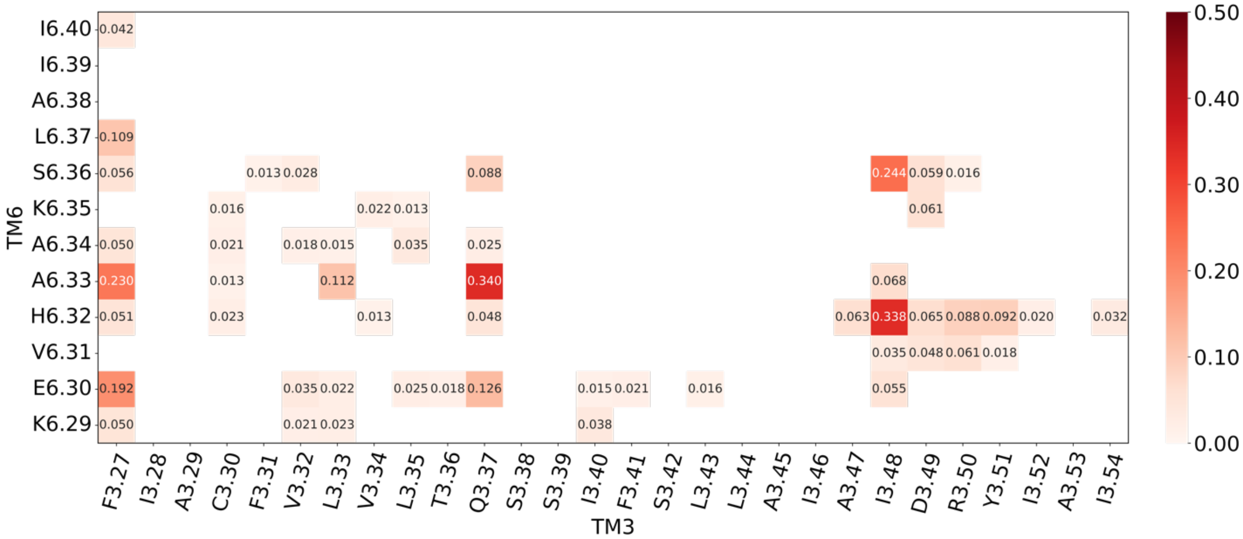


**Fig. S13. SHAP value heat map of residue pairs (Cα–Cα distances) indicating their contributions to distinguish clusters a and i.** Values are the average absolute SHAP values across five cross-validation folds.

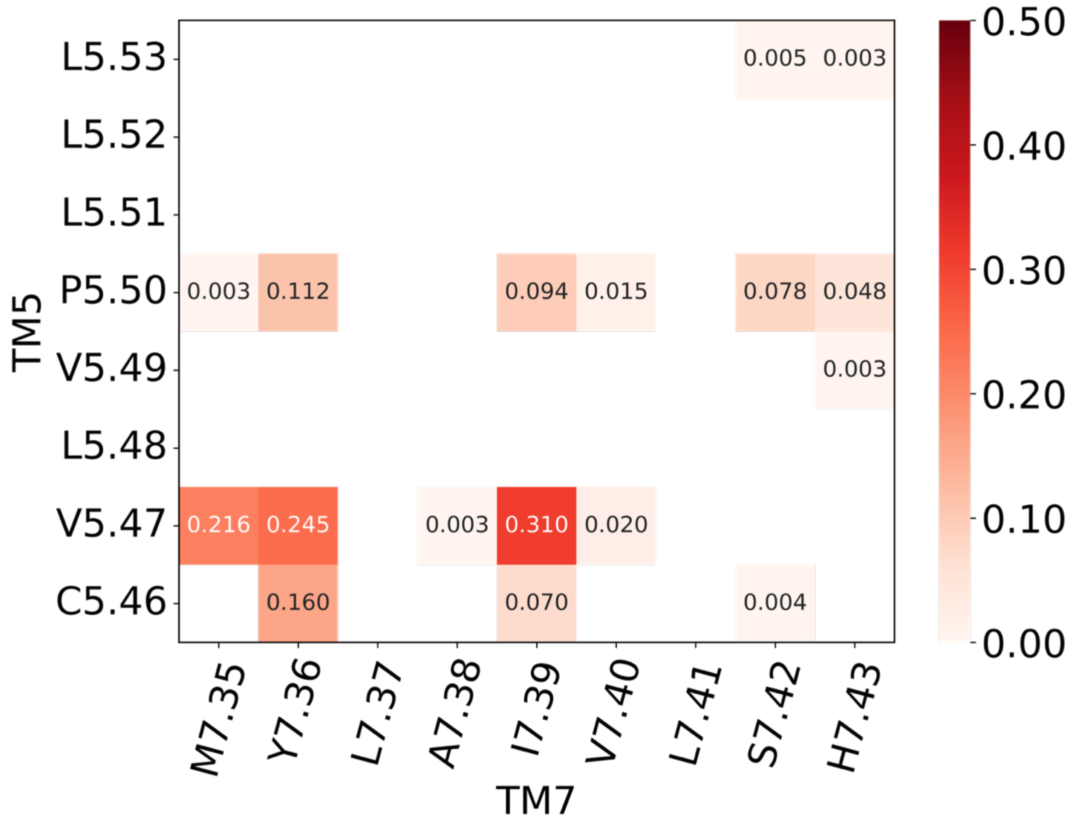


**Fig. S14. SHAP value heat map of residue pairs (Cα–Cα distances) indicating their contributions to distinguish clusters a and g.** Values are the average absolute SHAP values across five cross-validation folds.

**Table S1. Top 50 UMAP-HDBSCAN parameter settings ranked by DBCV scores**

| n_neighbors | min_dist | n_components | min_cluster_size | min_samples | #cluster | DBCV score |
|---|---|---|---|---|---|---|
| 500 | 1 | 9 | 1000 | 10 | 8 | 0.838337398 |
| 300 | 1 | 4 | 100 | 10 | 9 | 0.820977236 |
| 300 | 1 | 4 | 1000 | 10 | 7 | 0.805513416 |
| 100 | 1 | 10 | 100 | 10 | 7 | 0.785635528 |
| 100 | 1 | 10 | 300 | 10 | 7 | 0.785635528 |
| 100 | 1 | 10 | 500 | 10 | 7 | 0.785635528 |
| 100 | 1 | 10 | 700 | 10 | 7 | 0.785635528 |
| 100 | 1 | 10 | 1000 | 10 | 7 | 0.785635528 |
| 300 | 1 | 8 | 100 | 10 | 10 | 0.7836511 |
| 300 | 1 | 5 | 100 | 10 | 8 | 0.779840989 |
| 300 | 1 | 5 | 300 | 10 | 8 | 0.779840989 |
| 300 | 1 | 5 | 1000 | 10 | 8 | 0.779840989 |
| 500 | 1 | 9 | 300 | 10 | 9 | 0.76551831 |
| 500 | 1 | 9 | 500 | 10 | 9 | 0.76551831 |
| 500 | 1 | 9 | 700 | 10 | 9 | 0.76551831 |
| 200 | 1 | 5 | 100 | 10 | 8 | 0.761650781 |
| 200 | 1 | 5 | 300 | 10 | 8 | 0.761650781 |
| 200 | 1 | 5 | 500 | 10 | 8 | 0.761650781 |
| 200 | 1 | 5 | 700 | 10 | 8 | 0.761650781 |
| 200 | 1 | 5 | 1000 | 10 | 8 | 0.761650781 |
| 500 | 1 | 9 | 1000 | 50 | 8 | 0.760750647 |
| 300 | 1 | 4 | 300 | 10 | 8 | 0.759539547 |
| 300 | 1 | 4 | 500 | 10 | 8 | 0.759539547 |
| 300 | 1 | 4 | 700 | 10 | 8 | 0.759539547 |
| 500 | 1 | 9 | 100 | 10 | 10 | 0.756667153 |
| 100 | 1 | 6 | 100 | 10 | 7 | 0.7565576 |
| 100 | 1 | 6 | 300 | 10 | 7 | 0.7565576 |
| 100 | 1 | 6 | 500 | 10 | 7 | 0.7565576 |
| 100 | 1 | 6 | 700 | 10 | 7 | 0.7565576 |
| 100 | 1 | 6 | 1000 | 10 | 7 | 0.7565576 |
| 100 | 1 | 5 | 300 | 10 | 7 | 0.755438458 |
| 100 | 1 | 5 | 500 | 10 | 7 | 0.755438458 |
| 100 | 1 | 5 | 700 | 10 | 7 | 0.755438458 |
| 100 | 1 | 5 | 1000 | 10 | 7 | 0.755438458 |
| 500 | 1 | 9 | 100 | 50 | 10 | 0.752314476 |
| 100 | 1 | 7 | 100 | 10 | 7 | 0.746158673 |
| 100 | 1 | 7 | 300 | 10 | 7 | 0.746158673 |
| 100 | 1 | 7 | 500 | 10 | 7 | 0.746158673 |
| 100 | 1 | 7 | 700 | 10 | 7 | 0.746158673 |
| 100 | 1 | 7 | 1000 | 10 | 7 | 0.746158673 |
| 300 | 1 | 10 | 100 | 10 | 10 | 0.742842701 |

| | | | | | | |
|---|---|---|---|---|---|---|
| 300 | 1 | 10 | 300 | 10 | 9 | 0.740890321 |
| 300 | 1 | 10 | 500 | 10 | 9 | 0.740890321 |
| 300 | 1 | 10 | 700 | 10 | 9 | 0.740890321 |
| 300 | 1 | 10 | 1000 | 10 | 9 | 0.740890321 |
| 200 | 1 | 8 | 100 | 10 | 8 | 0.739678704 |
| 200 | 1 | 8 | 300 | 10 | 8 | 0.739678704 |
| 200 | 1 | 8 | 500 | 10 | 8 | 0.739678704 |
| 200 | 1 | 8 | 700 | 10 | 8 | 0.739678704 |
| 200 | 1 | 8 | 1000 | 10 | 8 | 0.739678704 |

**Table S2. Clusters b/j classification evaluation for model assessment**

| Fold | Accuracy_score | F1_score | Precision_score | Recall_score |
|---|---|---|---|---|
| 1 | 0.9950 | 0.9950 | 0.9950 | 0.9950 |
| 2 | 0.9950 | 0.9950 | 0.9950 | 0.9950 |
| 3 | 0.9975 | 0.9975 | 0.9975 | 0.9975 |
| 4 | 1.0000 | 1.0000 | 1.0000 | 1.0000 |
| 5 | 1.0000 | 1.0000 | 1.0000 | 1.0000 |

**Table S3. Clusters a/b classification evaluation for model assessment**

| Fold | Accuracy_score | F1_score | Precision_score | Recall_score |
|---|---|---|---|---|
| 1 | 1.0000 | 1.0000 | 1.0000 | 1.0000 |
| 2 | 1.0000 | 1.0000 | 1.0000 | 1.0000 |
| 3 | 1.0000 | 1.0000 | 1.0000 | 1.0000 |
| 4 | 1.0000 | 1.0000 | 1.0000 | 1.0000 |
| 5 | 1.0000 | 1.0000 | 1.0000 | 1.0000 |

**Table S4. Clusters b/c classification evaluation for model assessment**

| Fold | Accuracy_score | F1_score | Precision_score | Recall_score |
|---|---|---|---|---|
| 1 | 1.0000 | 1.0000 | 1.0000 | 1.0000 |
| 2 | 1.0000 | 1.0000 | 1.0000 | 1.0000 |
| 3 | 1.0000 | 1.0000 | 1.0000 | 1.0000 |
| 4 | 1.0000 | 1.0000 | 1.0000 | 1.0000 |
| 5 | 1.0000 | 1.0000 | 1.0000 | 1.0000 |

**Table S5. Clusters a/i classification evaluation for model assessment**

| Fold | Accuracy_score | F1_score | Precision_score | Recall_score |
|---|---|---|---|---|
| 1 | 0.9989 | 0.9989 | 0.9989 | 0.9989 |
| 2 | 0.9996 | 0.9996 | 0.9996 | 0.9996 |
| 3 | 1.0000 | 1.0000 | 1.0000 | 1.0000 |
| 4 | 0.9996 | 0.9996 | 0.9996 | 0.9996 |
| 5 | 1.0000 | 1.0000 | 1.0000 | 1.0000 |

**Table S6. Clusters a/f classification evaluation for model assessment**

| Fold | Accuracy_score | F1_score | Precision_score | Recall_score |
|---|---|---|---|---|
| 1 | 1.0000 | 1.0000 | 1.0000 | 1.0000 |
| 2 | 1.0000 | 1.0000 | 1.0000 | 1.0000 |
| 3 | 0.9995 | 0.9995 | 0.9995 | 0.9995 |
| 4 | 1.0000 | 1.0000 | 1.0000 | 1.0000 |
| 5 | 1.0000 | 1.0000 | 1.0000 | 1.0000 |

**Table S7. Clusters a/g classification evaluation for model assessment**

| Fold | Accuracy_score | F1_score | Precision_score | Recall_score |
|---|---|---|---|---|
| 1 | 1.0000 | 1.0000 | 1.0000 | 1.0000 |
| 2 | 1.0000 | 1.0000 | 1.0000 | 1.0000 |
| 3 | 1.0000 | 1.0000 | 1.0000 | 1.0000 |
| 4 | 1.0000 | 1.0000 | 1.0000 | 1.0000 |
| 5 | 0.9996 | 0.9996 | 0.9996 | 0.9996 |

**Table S8. Clusters g/h classification evaluation for model assessment**

| Fold | Accuracy_score | F1_score | Precision_score | Recall_score |
|---|---|---|---|---|
| 1 | 1.0000 | 1.0000 | 1.0000 | 1.0000 |
| 2 | 1.0000 | 1.0000 | 1.0000 | 1.0000 |
| 3 | 1.0000 | 1.0000 | 1.0000 | 1.0000 |
| 4 | 1.0000 | 1.0000 | 1.0000 | 1.0000 |
| 5 | 1.0000 | 1.0000 | 1.0000 | 1.0000 |

**Table S9. Number of lipids in the asymmetric membrane model (MM) system.** The value in the parentheses is the mole fraction % in each leaflet. Abbreviations are as follows: PC phosphatidylcholine; PE phosphatidyl ethanolamine; PS phosphatidyl serine; PSM 16:1,16:0 sphingomyelin; LSM 18:1,24:0 sphingomyelin; NSM 18:1,24:1 sphingomyelin; PAPC 16:0,20:4 PC; SOPC 18:0,18:1 PC; PLPC 16:0,18:2 PC; POPC 16:0,18:1 PC; OAPE 18:1,20:4 PE; PDPE 16:0,22:6 PE; PLQS 18:0,22:4 plasmalogen PE; PAPS 16:0,20:4 PS; CHOL cholesterol.

| Lipid class | Abbreviation | Asymmetric MM | |
|---|---|---|---|
| | | **Exo** | **Cyto** |
| SM | LSM | 48 (6.6) | 0 |
| | NSM | 56 (7.7) | 0 |
| | PSM | 72 (9.9) | 0 |
| PC | PAPC | 24 (3.3) | 0 |
| | SOPC | 40 (5.5) | 0 |
| | PLPC | 80 (11.0) | 80 (15.0) |
| | POPC | 0 | 32 (6.0) |
| PE | OAPE | 0 | 32 (6.0) |
| | PDPE | 0 | 72 (13.5) |
| PLAS | PLQS | 0 | 80 (15.0) |
| PS | PAPS | 0 | 120 (22.6) |
| CHOL | CHOL | 404 (55.8) | 116 (21.8) |

**Table S10. The receptor residues involved in the feature calculations.** RN: Residue Name; ID: Residue ID; BW: Ballesteros–Weinstein numbering.

| TM1 | | | TM2 | | | TM3 | | | TM4 | | | TM5 | | | TM6 | | | TM7 | | |
|---|---|---|---|---|---|---|---|---|---|---|---|---|---|---|---|---|---|---|---|---|
| **RN** | **ID** | **BW** | **RN** | **ID** | **BW** | **RN** | **ID** | **BW** | **RN** | **ID** | **BW** | **RN** | **ID** | **BW** | **RN** | **ID** | **BW** | **RN** | **ID** | **BW** |
| I | 10 | 1.36 | V | 40 | 2.38 | C | 77 | 3.25 | G | 118 | 4.39 | M | 177 | 5.38 | K | 227 | 6.29 | L | 269 | 7.34 |
| T | 11 | 1.37 | T | 41 | 2.39 | L | 78 | 3.26 | T | 119 | 4.40 | V | 178 | 5.39 | E | 228 | 6.30 | M | 270 | 7.35 |
| V | 12 | 1.38 | N | 42 | 2.40 | F | 79 | 3.27 | R | 120 | 4.41 | Y | 179 | 5.40 | V | 229 | 6.31 | Y | 271 | 7.36 |
| E | 13 | 1.39 | Y | 43 | 2.41 | I | 80 | 3.28 | A | 121 | 4.42 | F | 180 | 5.41 | H | 230 | 6.32 | L | 272 | 7.37 |
| L | 14 | 1.40 | F | 44 | 2.42 | A | 81 | 3.29 | K | 122 | 4.43 | N | 181 | 5.42 | A | 231 | 6.33 | A | 273 | 7.38 |
| A | 15 | 1.41 | V | 45 | 2.43 | C | 82 | 3.30 | G | 123 | 4.44 | F | 182 | 5.43 | A | 232 | 6.34 | I | 274 | 7.39 |
| I | 16 | 1.42 | V | 46 | 2.44 | F | 83 | 3.31 | I | 124 | 4.45 | F | 183 | 5.44 | K | 233 | 6.35 | V | 275 | 7.40 |
| A | 17 | 1.43 | S | 47 | 2.45 | V | 84 | 3.32 | I | 125 | 4.46 | A | 184 | 5.45 | S | 234 | 6.36 | L | 276 | 7.41 |
| V | 18 | 1.44 | L | 48 | 2.46 | L | 85 | 3.33 | A | 126 | 4.47 | C | 185 | 5.46 | L | 235 | 6.37 | S | 277 | 7.42 |
| L | 19 | 1.45 | A | 49 | 2.47 | V | 86 | 3.34 | I | 127 | 4.48 | V | 186 | 5.47 | A | 236 | 6.38 | H | 278 | 7.43 |
| A | 20 | 1.46 | A | 50 | 2.48 | L | 87 | 3.35 | C | 128 | 4.49 | L | 187 | 5.48 | I | 237 | 6.39 | T | 279 | 7.44 |
| I | 21 | 1.47 | A | 51 | 2.49 | T | 88 | 3.36 | **W** | **129** | **4.50** | V | 188 | 5.49 | I | 238 | 6.40 | N | 280 | 7.45 |
| L | 22 | 1.48 | **D** | **52** | **2.50** | Q | 89 | 3.37 | V | 130 | 4.51 | **P** | **189** | **5.50** | V | 239 | 6.41 | S | 281 | 7.46 |
| G | 23 | 1.49 | I | 53 | 2.51 | S | 90 | 3.38 | L | 131 | 4.52 | L | 190 | 5.51 | G | 240 | 6.42 | V | 282 | 7.47 |
| **N** | **24** | **1.50** | A | 54 | 2.52 | S | 91 | 3.39 | S | 132 | 4.53 | L | 191 | 5.52 | L | 241 | 6.43 | V | 283 | 7.48 |
| V | 25 | 1.51 | V | 55 | 2.53 | I | 92 | 3.40 | F | 133 | 4.54 | L | 192 | 5.53 | F | 242 | 6.44 | N | 284 | 7.49 |
| L | 26 | 1.52 | G | 56 | 2.54 | F | 93 | 3.41 | A | 134 | 4.55 | M | 193 | 5.54 | A | 243 | 6.45 | **P** | **285** | **7.50** |
| V | 27 | 1.53 | V | 57 | 2.55 | S | 94 | 3.42 | I | 135 | 4.56 | L | 194 | 5.55 | L | 244 | 6.46 | F | 286 | 7.51 |
| C | 28 | 1.54 | L | 58 | 2.56 | L | 95 | 3.43 | G | 136 | 4.57 | G | 195 | 5.56 | C | 245 | 6.47 | I | 287 | 7.52 |
| W | 29 | 1.55 | A | 59 | 2.57 | L | 96 | 3.44 | L | 137 | 4.58 | V | 196 | 5.57 | W | 246 | 6.48 | Y | 288 | 7.53 |
| A | 30 | 1.56 | I | 60 | 2.58 | A | 97 | 3.45 | T | 138 | 4.59 | Y | 197 | 5.58 | L | 247 | 6.49 | A | 289 | 7.54 |
| V | 31 | 1.57 | P | 61 | 2.59 | I | 98 | 3.46 | P | 139 | 4.60 | L | 198 | 5.59 | **P** | **248** | **6.50** | Y | 290 | 7.55 |
| W | 32 | 1.58 | F | 62 | 2.60 | A | 99 | 3.47 | | | | R | 199 | 5.60 | L | 249 | 6.51 | R | 291 | 7.56 |
| L | 33 | 1.59 | A | 63 | 2.61 | I | 100 | 3.48 | | | | I | 200 | 5.61 | H | 250 | 6.52 | I | 292 | 7.57 |
| | | | I | 64 | 2.62 | D | 101 | 3.49 | | | | F | 201 | 5.62 | I | 251 | 6.53 | R | 293 | 7.58 |
| | | | T | 65 | 2.63 | **R** | **102** | **3.50** | | | | L | 202 | 5.63 | I | 252 | 6.54 | E | 294 | 7.59 |
| | | | | | | Y | 103 | 3.51 | | | | A | 203 | 5.64 | N | 253 | 6.55 | F | 295 | 7.60 |
| | | | | | | I | 104 | 3.52 | | | | A | 204 | 5.65 | C | 254 | 6.56 | R | 296 | 7.61 |
| | | | | | | A | 105 | 3.53 | | | | R | 205 | 5.66 | F | 255 | 6.57 | | | |
| | | | | | | I | 106 | 3.54 | | | | R | 206 | 5.67 | | | | | | |
| | | | | | | | | | | | | Q | 207 | 5.68 | | | | | | |
| | | | | | | | | | | | | L | 208 | 5.69 | | | | | | |
| | | | | | | | | | | | | K | 209 | 5.70 | | | | | | |
| | | | | | | | | | | | | Q | 210 | 5.71 | | | | | | |

**Movie S1. HDBSCAN clustering on the UMAP projection.** This video shows distinct conformational clusters across nine simulated systems in three UMAP dimensions. It supplements Fig. 1.

**Movie S2. HDBSCAN clustering on the UMAP embedding including six class A arrestin-bound receptors represented by diamonds.** This video shows that the arrestin coupled structures stay in the boundary between the fully active conformations (cluster a, in green) and the pseudo-active state conformations (cluster i, in tan) in three UMAP dimensions. It supplements Fig. 5.

The movies S1 and S2 are available on https://doi.org/10.5281/zenodo.19161825.
